# Defining state-selective lipid binding to brain GPCRs - introducing REVEAL

**DOI:** 10.64898/2026.05.01.722320

**Authors:** Sophie A. S. Lawrence, Carla Kirschbaum, Corinne A. Lutomski, Haigang Song, Joshua D. Hinkle, Frances I. Butroid, Rafael Melani, John E. P. Syka, Christopher Mullen, Tarick J. El-Baba, Carol V. Robinson

## Abstract

Membrane lipids are central regulators of G protein–coupled receptor (GPCR) function. Defining receptor-specific lipid interactions in native, fully modified mammalian systems remains challenging. Extensive post-translational modifications generate heterogeneous proteoforms that confound conventional mass spectrometry approaches. Here we introduce REVEAL (REceptor enVironment Elucidation by Activated Lipid-release), an automated native top-down mass spectrometry strategy that discriminates specifically bound lipids from background. Applied here to intact, heterogeneous mammalian membrane protein complexes, incubated with a brain polar lipid extract (>1000 components), we define receptor-specific lipid-binding for two neuronal class C GPCRs. We show that agonism remodels lipid occupancy, selectively enriching a reduced repertoire of bound lipids. Plasmalogen lipids emerge as persistent binders across all conformational states of the metabotropic glycine receptor and are preferentially depleted under oxidative stress, implying a protective role at the receptor surface. These findings position lipids as dynamic regulators of both function and response to the cellular redox environment.

## Introduction

A central challenge in studying binding to mammalian membrane proteinsis their intrinsic molecular heterogeneity. Proteoform diversity for G protein-coupled receptors (GPCRs), solute carriers (SLCs), and ion channels arises from extensive post-translational modifications (PTM), including glycosylation, phosphorylation and lipidation. This complex ensemble of proteoforms overlap in mass, and the resulting heterogeneity obscures direct detection of individual ligand binding events. Most experimental strategies consequently rely on simplified systems such as engineered proteins or deglycosylated receptors. However, these approaches may overlook biologically relevant interactions present in native contexts ^1-3^.

The application of native mass spectrometry (nMS) for detecting non-covalent protein-ligand interactions by tracking changes in mass is straightforward in theory ^4-6^. In practice however, applying such assays to mammalian membrane proteins is complicated by proteoform heterogeneity which confounds discrimination between ligand-free and ligand-bound states. Native top-down MS (nTDMS) partially addresses this limitation by isolating and activating precursor ions to yield product ion spectra reporting on both the composition and structure of protein-ligand complexes ^7-10^. nTDMS is not able to resolve overlapping peaks arising from bound and unbound states in the mass spectra of large, heavily modified proteins. We previously approached this problem through a native MS-guided lipidomics workflow ^3^; however, this strategy still cannot distinguish specifically bound lipids from those loosely associated or present in empty micelles.

To define direct ligand-binding events across different conditions we introduce REVEAL (REceptor enVironment Elucidation by Activated Ligand-release). REVEAL is an automated untargeted nTDMS strategy for direct identification of ligands specifically bound to fully modified mammalian membrane proteins. Analogous to data-independent acquisition in bottom-up proteomics ^11-14^, REVEAL sequentially interrogates non-overlapping narrow *m*/*z* windows across the mass spectrum. All precursor ions within each window are isolated and activated simultaneously to release bound ligands/lipids within the mass spectrometer. Comparison of the resulting product ion spectra against the intact precursor distribution (MS^1^) enables specifically bound lipids to be distinguished from those non-specifically co-isolated within the background micelle environment.

Given the importance of lipids in shaping GPCR structure and function ^15-17^ we apply REVEAL to define lipid binding preferences for the metabotropic glycine receptor (mGlyR) and the metabotropic glutamate receptor 2 (mGluR2). These structurally distinct, dimeric class C GPCRs are highly expressed in the human brain and represent promising therapeutic targets for neurological and psychiatric disorders ^18-23^. Structural studies of both receptors have identified lipid-like electron densities within their transmembrane (TM) domains, implicating specific lipids in the regulation of receptor structure and activation ^24-30^. Nevertheless, extensive N-glycosylation and other PTMs have hindered direct identification of individual lipid–binding events.

Here, following expression and purification of these two receptors from human cell lines we incubated them with brain lipid extracts, comprised of >1,500 distinct lipids. We then applied REVEAL to define receptor-specific lipid-binding fingerprints which comprise distinct classes of phospholipids and sphingolipids. Agonism of both receptors induces conformational rearrangements at transmembrane interfaces that coincide with a broad remodelling of the lipid-binding landscape. A small subset of lipids susceptible to oxidative stress bound to mGlyR regardless of conformation ^31-34^. Consistent with this subset of lipids serving a regulatory role in mGlyR function, exposure to oxidative stress selectively disrupted the mGlyR-plasmalogen interactions. Moreover, loss of these plasmalogen lipids potentiated the basal-level activity of the receptor. Given that production of stress-induced reactive oxygen species (ROS) is correlated with upregulation of mGlyR stress-response signalling in brain ^23,27,28^, our findings suggest a lipid-mediated route by which oxidative stress may modulate receptor activity.

## Results

### REVEAL defines direct lipid-binding repertoire of mGlyR

To overcome the challenges associated with studying GPCR-lipid interactions, we developed REVEAL to interrogate associated ligands within intact micellated receptor complexes (**Fig. 1a**). Where individual peaks corresponding to receptor-ligand binding cannot be resolved in a native spectrum, REVEAL links direct binding events to the receptor by aligning product ion chromatograms (from MS^2^) with nMS distributions. For associated lipids that bind non-specifically, poor alignment between their intensity distributions and those of the intact receptor, enables us to classify them as originating from background micelles.

**Figure 1.**
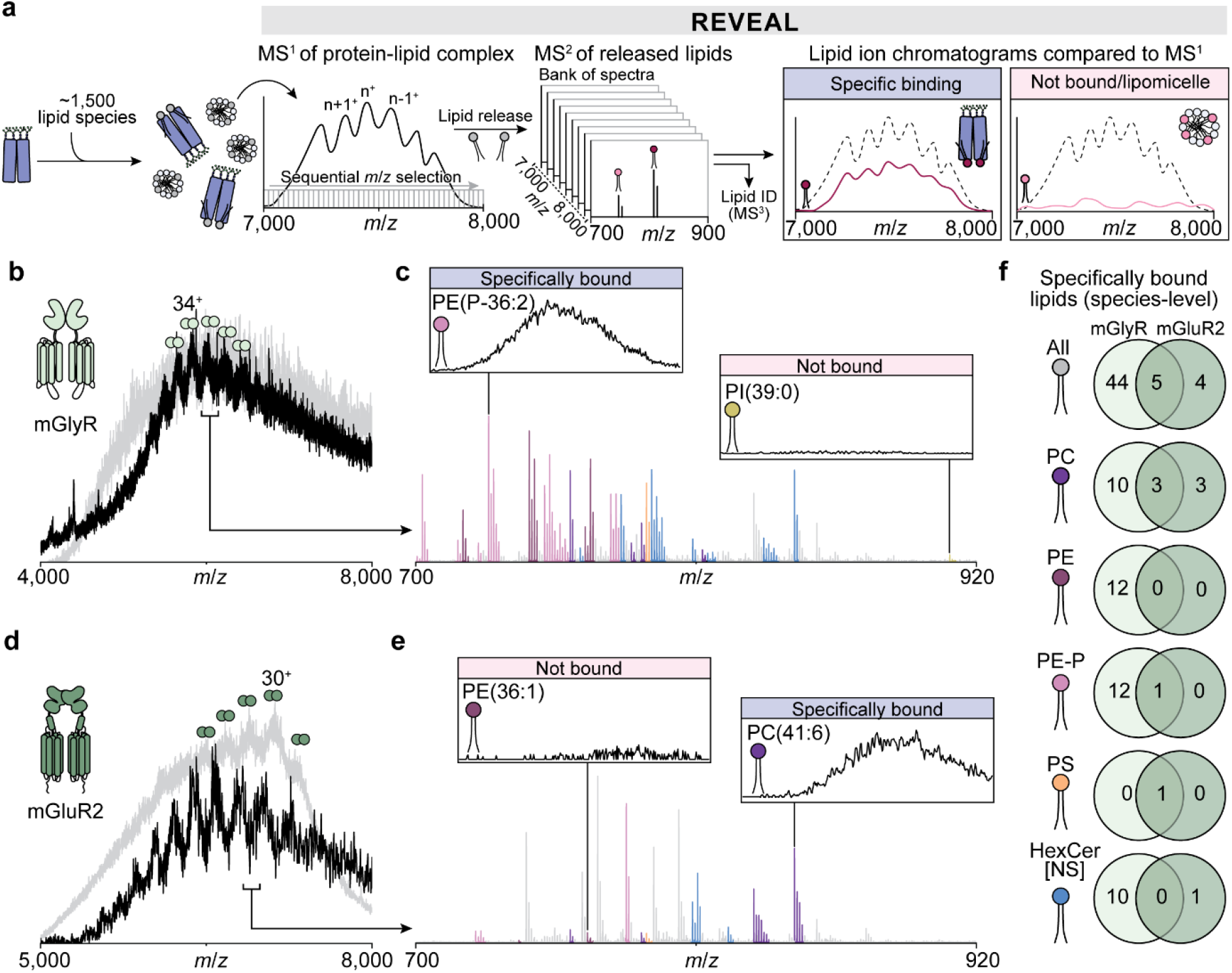
Uncovering unique lipid-binding preferences of Class-C GPCRs using REVEAL. (**a**) Overview of the REVEAL (REceptor enVironment Elucidation by Activated Lipid-release) workflow to discriminate specifically bound from co-isolating lipids. Briefly, a protein is incubated with a large repertoire of ligands/lipids before a nMS/MS^1^ spectrum is recorded. Regions are then selected sequentially across the MS^1^ spectrum using discrete, nonoverlapping windows; all precursors within each window are dissociated, and the resulting MS^2^ product ion spectra are recorded. Extracted ion chromatograms are generated for each product ion of interest to yield time-domain data which is converted into the *m*/*z* domain using bins dictated by the window size. These product ion spectra are overlaid against the MS^1^ spectrum. Product ions whose intensity distributions overlap with the MS^1^ spectrum are classified as likely binders; those with intensity distributions in different *m*/*z* regions or distributed across the entire *m*/*z* range are classified as co-isolated but not directly binding. (**b**) Native mass spectrum of dimeric delipidated mGlyR (black) overlaid with that of mGlyR incubated with a 10-fold molar excess of a brain lipid extract (grey). The measured mass of dimeric mGlyR is 192,075 ± 161 Da. (**c**) Low *m*/*z* region of the MS^2^ spectrum following nTDMS of mGlyR (quadrupole selection at *m*/*z* 6000, 100 *m*/*z* window, 120 V HCD). Lipid identities were assigned using ion-trap MS^3^ (**Supplementary Figs. 2-7**) and database searching; fragmentation patterns indicate lipid class, while multiple diagnostic fragments in MS^3^ spectra indicate co-isolated precursors. Lipids are coloured by class. Inset: REVEAL distinguished specifically bound from bulk lipids (**Supplementary Figs. 9-10**). Reconstructed precursor spectra for PE(P-36:2) (*m*/*z* 728.5; bound) and PI(39:0) (*m/z* 909.5; not bound) are shown. (**d**) Native mass spectrum of dimeric delipidated mGluR2 (black) overlaid onto that of mGluR2 incubated with a 10-fold molar excess of a brain lipid extract (grey). The measured mass of mGluR2 is 210,808 ± 206 (black) and lipid-bound mGluR2 is 213,473 ± 240 Da (grey). (**e**) Low *m*/*z* region of the MS^2^ spectrum following nTDMS of mGluR2 (quadrupole selection at *m*/*z* 7000, 100 *m*/*z* window, 120 V HCD). Lipids were assigned by ion-trap MS^3^ (**Supplementary Figs. 2-7**) and database searching, as in **c**. Inset: REVEAL distinguished specifically bound from bulk lipids (**Supplementary Figs. 9**,**12**); reconstructed precursor spectra for PE(36:1) (*m/z* 788.6; not bound) and PC(46:1) (*m/z* 848.6; bound) are shown. (**f**) Venn diagrams summarising lipid binding to mGlyR (pale green) and mGluR2 (dark green) at the species level (a sum composition of carbons and double bonds in the fatty acid chains); separated by lipid class.

To characterise the repertoire of brain lipids that associate with mGlyR, we first recorded a native mass spectrum of mGlyR following extensive delipidation. Broad but resolved charge states were observed centred near *m*/*z* 6000 (**Fig. 1b**), corresponding to a complex of mass 192,075 ± 161 Da (cf. dimeric mGlyR, 191,535 Da) (**Supplementary Table 1**). The breadth of the charge state peaks reflects the coexistence of multiple mGlyR proteoforms arising from glycosylation, palmitoylation, or other PTMs. Following delipidation we challenged mGlyR with a brain polar lipid extract, which comprises over 1000 unique polar lipid species (**Supplementary Fig. 1**). The resulting native mass spectrum **(Fig. 1b)** remained centred around *m*/*z* 6000, but the charge state peaks became unresolved, a hallmark of extensive lipid binding.

To broadly identify of the associated lipids, we released all lipids from a broad *m*/*z* range and recorded the resulting product ion mass spectrum (MS^2^). Over 60 abundant, well-resolved peaks were observed (**Fig. 1c**). These peaks were assigned to distinct lipids via their characteristic fragmentation patterns (MS^3^) (**Supplementary Table 2**). In total, we identified the following lipid species: 19 phosphatidylcholines (PC), 32 phosphatidylethanolamines (PE), one phosphatidylinositol (PI) and one phosphatidylserine (PS) (**Supplementary Fig. 2-5**). Structural analysis of *sn*-1 and *sn*-2 fragment ion masses revealed that a substantial proportion of the PE lipids were of the plasmalogen form (PE-P), containing a distinct vinyl ether linkage at the s*n*-1 position (**Supplementary Fig. 6, Supplementary Table 3-4**) ^35,36^. Although lipids from nearly all major classes were captured, cholesterol, a well-established membrane protein binding lipid ^27,37-39^, was not detected consistent with its poor ionisation efficiency ^40^. Beyond these glycerophospholipids, 12 sphingosine hexosylceramides (HexCer[NS]) were also identified (**Supplementary Fig. 7**) ^41^. Along with plasmalogens, HexCer[NS] species are known to play important roles in organising and regulating signalling processes in neuronal membranes ^42,43^.

**Figure 2.**
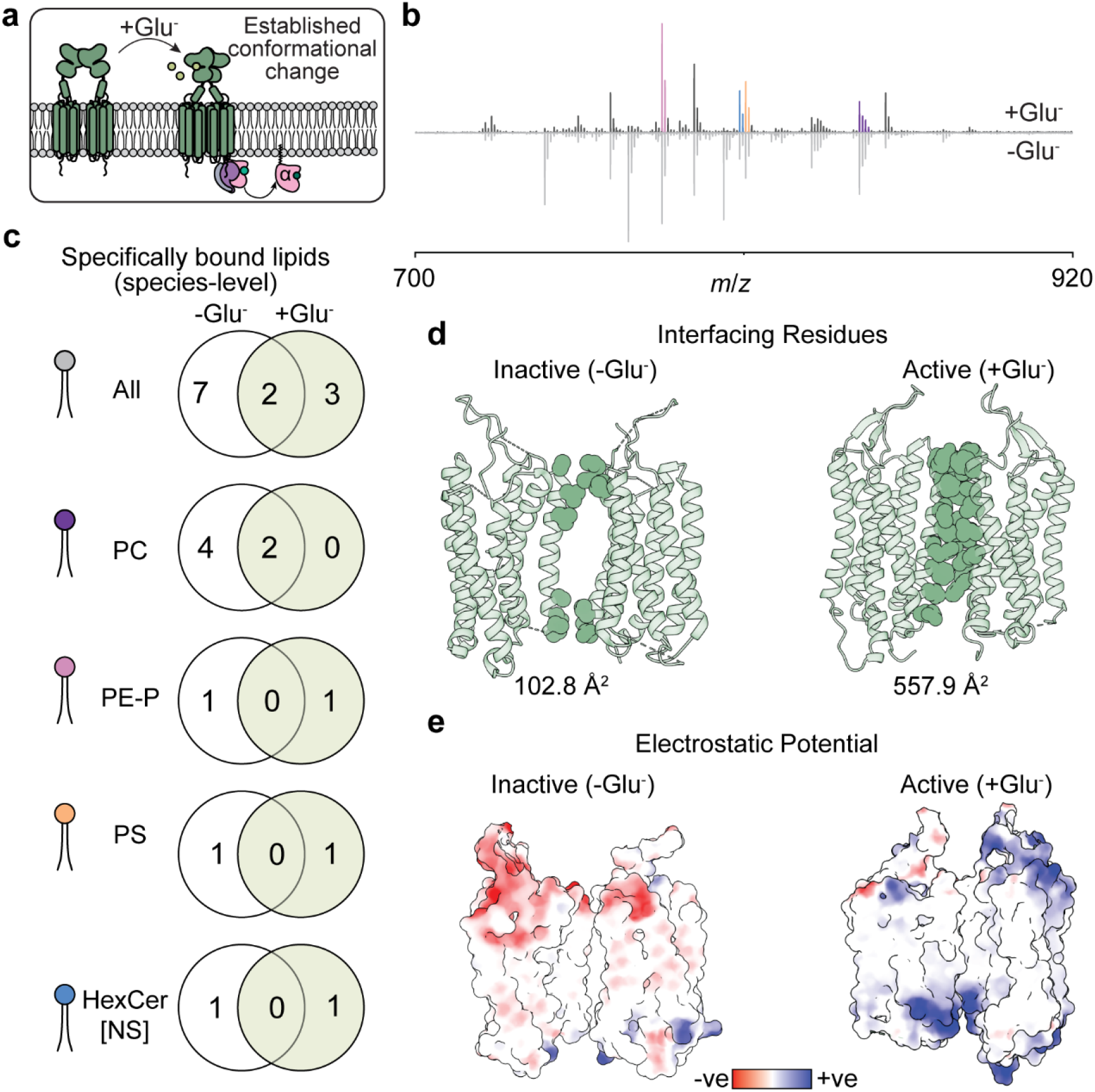
Glutamate agonism of mGluR2 restricts the lipid-binding repertoire. (**a**) Cartoon representation of the impact of glutamate on mGluR2 conformation and downstream signalling pathways. (**b**) Low *m*/*z* region of the MS^2^ spectrum from nTDMS of mGluR2 incubated with a 10-fold molar excess of brain lipids in the presence (top) or absence (bottom) of glutamate (quadrupole selection at *m*/*z* 7500, 1000 *m*/*z* window; 120 V HCD). Isotopic envelopes corresponding to lipids released during this process are shown. Classification of the lipids was confirmed by ion-trap MS^3^ (**Supplementary Figs. 2-7**), and chain level lengths were assigned by database search. Specifically bound lipids were determined by REVEAL (**Supplementary Fig. 13**) and are coloured by class. (**c**) Venn diagrams showing the number of lipids specifically bound to mGluR2 with and without glutamate agonism, separated by lipid class. (**d**) Interfaces between the TM helices in inactive (PDB 7MTQ) and active (PDB 7MTS) mGluR2. Residues contributing to inter-protomer contacts within the transmembrane domain (defined using PDBePISA) are coloured; calculated interfacial surface areas are indicated. (**e**) Electrostatic potential surfaces of cryo-EM structures of inactive (PDB 7MTQ) and active (PDB 7MTS) mGluR2. Negative and positive electrostatic potential is indicated in red and blue, respectively.

We next applied REVEAL to discriminate between lipids specifically bound to mGlyR and those arising from association due to their co-isolation in micelles. By generating a bank of 400 individual MS^2^ spectra (**Supplementary Fig. 8**) and reconstructing the precursor ion distribution for each identified lipid, we classified each species as either specifically bound or associated (*viz*. originating from background micelles). Of the lipids identified, 49 were classified as specifically bound to mGlyR, while approximately 15 lipids were assigned to background (**Fig. 1f, Supplementary Fig. 9-10**). Multiple PE-P species were confirmed as specific binders, whereas PI lipids showed no coincidence between their intensities and the native spectrum of lipid-bound mGlyR. Considering the specific binders, we assessed whether mGlyR exhibits selectivity for particular PE-P chain lengths or degrees of unsaturation. To do this we compared the relative abundances of bound PE-P species with those in the brain lipid extract (**Supplementary Fig. 11**). We observed a broadly similar distribution of chain lengths. Our results suggest that mGlyR does not strongly discriminate among specifically bound PE-P variants on the basis of acyl chain length or degree of unsaturation.

### Divergent lipid-binding preferences across class C GPCRs

We next compared the lipid-binding preferences of mGlyR with mGluR2. We recorded a native mass spectrum of delipidated mGluR2 and observed a series of well-resolved charge state peaks corresponding to a molecular weight of 210,808 ± 206 Da, consistent with fully glycosylated dimeric mGluR2 (208,780 Da) (**Fig. 1d**). We then incubated mGluR2 with the brain polar lipid extract as above and recorded a mass spectrum (**Fig. 1d**). The resulting charge state distribution was broad and consisted of wide charge state peaks that were shifted to a higher *m*/*z* (deconvoluted mass of 213,473 ± 240 Da), indicative of lipid binding. Analysis of the MS^2^ spectrum of lipids released directly from mGluR2 (without our stepwise strategy of REVEAL) uncovered 18 associated lipids from the PC, PE-P, PS and HexCer[NS] families (**Fig. 1e, Supplementary Fig. 2-7**). Next by applying REVEAL only nine of these lipids were classified as specifically bound, including six PC, and individual PE-P, PS, and HexCer[NS] lipids (**Fig. 1f, Supplementary Fig. 8-9, 12**).

Comparison of the two receptors exhibited a substantially broader repertoire of specifically bound lipids (49 species for mGlyR compared with 9 species for mGluR2) (**Fig. 1f, Supplementary Fig. 9)**. Both receptors showed a preference for PC lipids. Of note, both PE and PE-P species were disproportionately represented among the specifically bound lipids in mGlyR relative to their abundance in neuronal membranes, indicative of selective enrichment of lipids by mGlyR. Both mGlyR and mGluR2 bound specifically to PS, an anionic inner-leaflet lipid that is inherently challenging to detect by positive-ion mode mass spectrometry ^44^. Considering the bilayer distributions of these lipid classes implies their likely binding locations. For example, outer-leaflet lipids such as PC and HexCer typically occupy the annular belt surrounding the receptor periphery. By contrast inner-leaflet lipids (PE, PE-P, and PS) are often located at inter-protomer interfaces and have been shown to directly stabilise membrane protein structure and function ^45^. This raises the possibility that these protein-lipid interactions may be influenced by receptor activation.

### Agonism streamlines the lipid repertoire of mGluR2

The stark differences in lipid-binding profiles between mGlyR and mGluR2 suggest that lipid interactions are shaped by the receptor and may change in response to different receptor conformations. Consistent with this, structural studies of mGluR2 have shown that agonist binding drives large-scale conformational rearrangements that expose distinct transmembrane surfaces ^24-26^. To explore this, we examined lipid binding to mGluR2 in the glutamate-bound state and compared it to the resting state of the protein (**Fig. 2a**). In this activated state, 30 associated lipids were assigned. Application of our REVEAL workflow subsequently reduced these to five lipids designated as specifically bound (**Fig. 2b-c**). This is a marked reduction from the nine lipids identified in the resting state. The five lipid species identified as specifically bound in the glutamate-bound state are PC, PE-P, HexCer[NS] and PS (**Fig. 2c, Supplementary Fig. 9, 13**).

We considered a structural basis to explain this selectivity in lipid binding focussing on activation-dependent changes at the TM dimer interface reported in cryo-EM structures ^24-26^. In the inactive state, the interface is asymmetric and dominated by TM3–TM4 contacts. Activation reorganises the interface symmetrically around the TM6–TM6 contacts increasing the protein-protein contact area approximately five-fold (**Fig. 2d**). Correspondingly, the solvent-accessible volume at the dimer interface decreases approximately three-fold upon activation, substantially reducing the capacity to accommodate a diverse lipid repertoire (**Supplementary Fig. 14**). Interhelical distances within individual protomers remain largely unchanged, indicating that lipid displacement is localised to the dimer interface rather than occurring globally across the receptor. Considering also changes in charge; the inactive receptor exhibits negatively and positively charged extracellular and intracellular-facing surfaces respectively. Upon activation this charge is redistributed to positive towards the central interfacial cavity (**Fig. 2e**). Together, consolidation of the dimer interface and redistribution of electrostatic surface potential provide a structural rationale for a reduced lipid repertoire upon activation. Accordingly, PC species associated with the extracellular leaflet are depleted upon activation. By contrast, select PE-P and PS species, localised to the inner leaflet, are retained. These findings support a model in which activation eliminates loosely associated lipids, through increased protein-protein contacts, while retaining a subset of bound lipids within structurally confined interfacial pockets.

### Lipid binding scales with receptor activation in mGlyR

Analogous studies of agonist-bound mGlyR have not yet been reported. It is established however that mGlyR signals through a non-canonical pathway. G_i/o_ activity is modulated by regulator of G protein signalling (RGS) proteins (notably RGS7) that accelerate GTP hydrolysis on G_α_ subunits to promote rapid signal termination ^27,30^. Glycine binding relieves this constraint, facilitating G_i/o_ signalling ^28^ and suggesting that mGlyR may sample distinct conformational ensembles to facilitate this signalling (**Fig. 3a**). Cryo-EM structures have been reported for apo mGlyR and for a nanobody-stabilised form exhibiting glycine-like activity ^27,29,30^. In the apo state, mGlyR adopts a non-canonical TM architecture, forming an inverted V-shaped helical arrangement with tight TM4-TM5 contacts near the extracellular face and a pronounced intracellular opening. Relative to mGluR2, mGlyR has an increased inter-helical distance and distinct electrostatic features within the central cavity, both of which suggest an enhanced capacity for lipid interdigitation between protomers (**Supplementary Fig. 15**).

**Figure 3.**
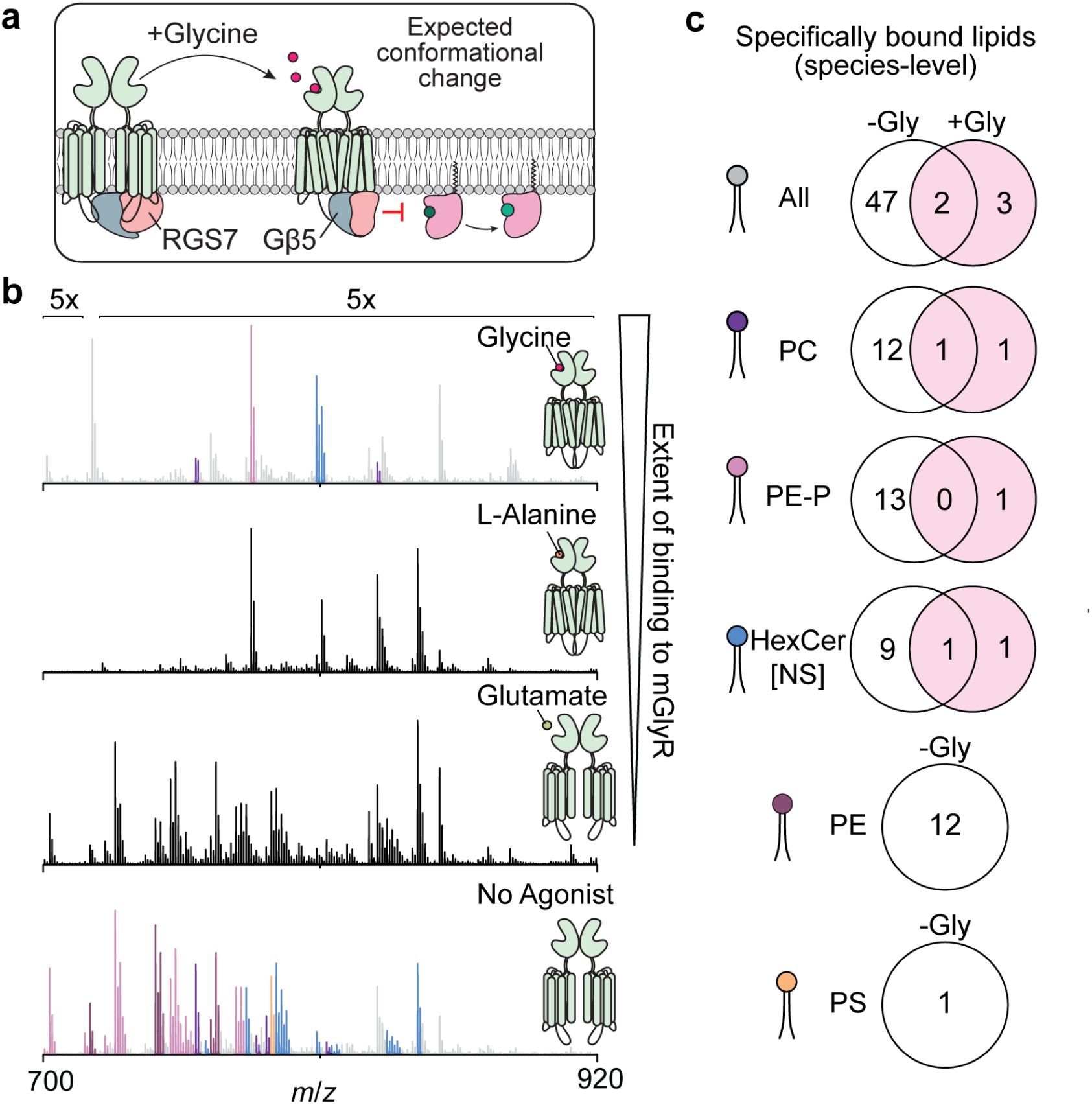
Agonism of mGlyR narrows the lipid-binding repertoire but preserves key lipid interactions. (**a**) Cartoon representation of the impact of glycine on mGlyR conformation and signalling pathways. (**b**) Low *m*/*z* region of MS^2^ spectrum of mGlyR incubated with a 10-fold molar excess of brain lipids in the presence of glycine (top), L-alanine (second from top), glutamate (second from bottom), no agonist (bottom) (quadrupole selection at *m*/*z* 7000, 1000 *m*/*z* wide, 120 V HCD activation). Isotopic envelopes corresponding to lipid adducts are observed. Lipids were confirmed by ion-trap MS^3^ (**Supplementary Figs. 2-7**). Side chain lengths of the lipid class were assigned by database search. For glycine, specifically bound lipids were determined by REVEAL (**Supplementary Fig. 16**) and are coloured by class. (**c**) Venn diagrams showing the number of specifically bound lipids to mGlyR with and without glycine agonism, separated by lipid class.

To determine whether lipid binding to mGlyR is conformation-dependent, we incubated delipidated mGlyR with brain polar lipids in the presence or absence of glycine. Using REVEAL, we found that following glycine agonism only five peaks were observed consistent with specifically bound lipids in this activated state (**Fig. 3b-c, Supplementary Fig. 16**). These lipids are assigned as PC, PE-P and HexCer[NS] species (**Fig. 3b, Supplementary Fig. 9**). This observation therefore represents a substantial narrowing of the lipid repertoire relative to the apo receptor. Only two species, PE(P-40:3) and a HexCer[NS] lipid, are common to activated mGluR2.

To further probe the relationship between receptor conformational state and lipid binding, we performed parallel experiments with glutamate, a non-agonist for mGlyR, and L-alanine, a structural analogue of glycine with weak partial agonist activity ^28^. Glutamate produced a lipid-binding profile indistinguishable from the agonist-free receptor, confirming that the observed changes in lipid repertoire are coupled to receptor agonism (**Fig. 3b**). In contrast, L-alanine yielded an intermediate binding profile, lying between the apo and fully agonised states, with binding observed across multiple lipid classes, including PE-P. This profile is consistent with a reduced lipid repertoire but without the reduction induced by full activation (**Fig. 3b**). This graded response demonstrates that the reduced lipid repertoire and increased lipid selectivity scales with the degree of receptor activation. Moreover, this observation provides evidence that lipid binding is coupled to conformational changes along the activation pathway.

The retention of long-chain PE(P-40:3) across both activated mGlyR and mGluR2 is particularly notable given its relatively low abundance in the brain lipid extract (**Supplementary Fig. 1, 11**). PE(P-40:3) encompasses multiple *sn*-1/*sn*-2 acyl chain combinations (**Supplementary Fig. 6**). As it is the sole lipid species preserved within two activated class C GPCRs, it implies that activation imposes stringent geometric and electrostatic constraints satisfied by PE(P-40:3) (e.g., hydrophobic length, unsaturation and headgroup orientation). Given the established link between mGlyR and stress-responsive signalling in the brain ^23,27,28,46^, the consistent enrichment of PE-P interactions regardless of activation state raises the possibility that these lipids are functionally important beyond receptor activation, such as conferring selective sensitivity of mGlyR to oxidative conditions. We investigate this hypothesis directly in the following section.

### Plasmalogens act as redox buffers against oxidative modification of mGlyR

Given the selective association of mGlyR with antioxidant plasmenyl-PE lipids, and its reported involvement in stress-related disorders ^23,27^, we next investigated whether oxidative stress alters mGlyR–lipid interactions. ROS damage both membrane proteins and lipids ^47-50^ (**Fig. 4a**), suggesting that oxidative modification of receptor-associated lipids may couple redox signals to changes in mGlyR structure and function.

**Figure 4.**
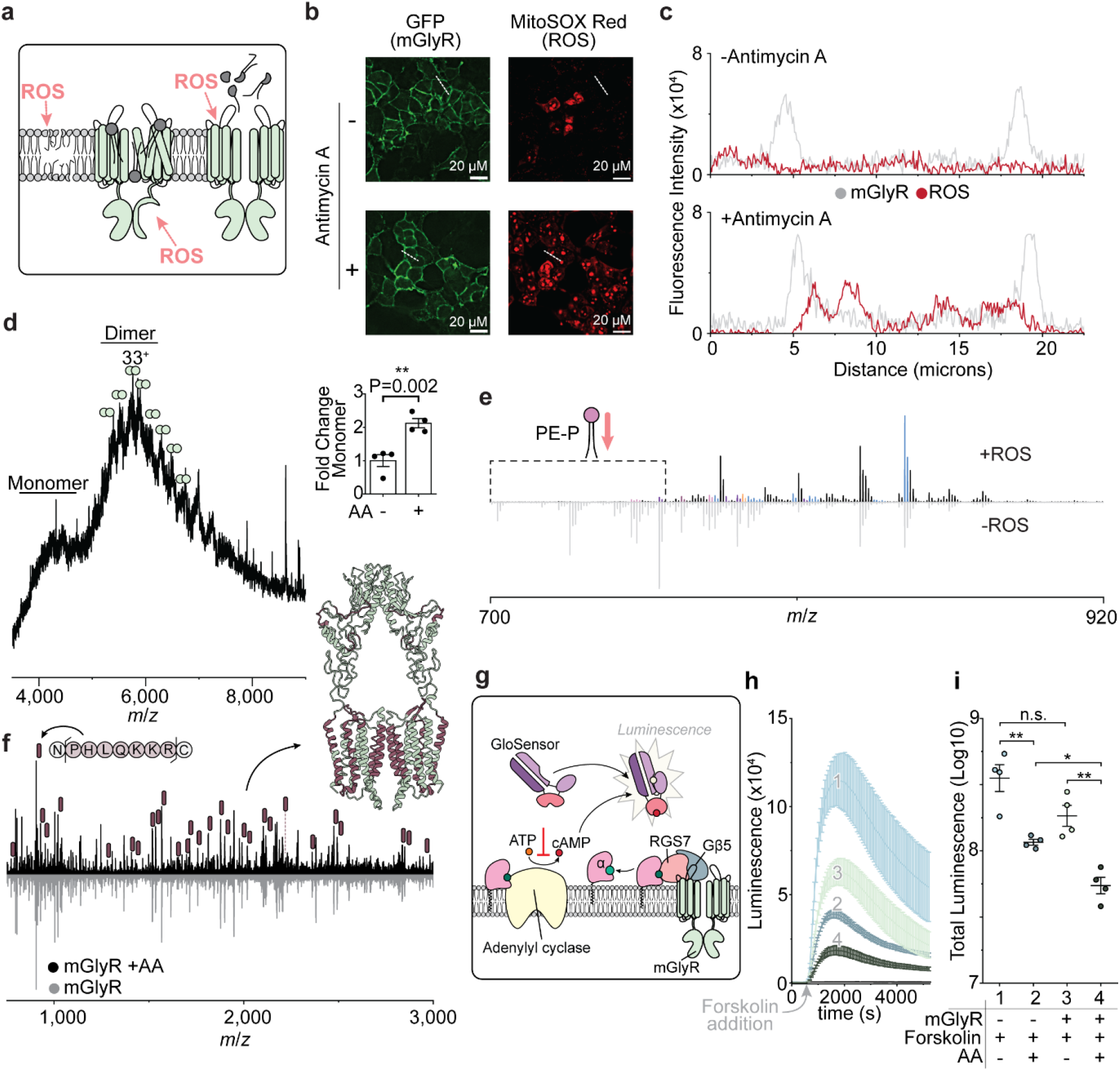
PE-Ps act as redox buffers against oxidative modification of mGlyR. (**a**) Cartoon representation of the putative impacts of ROS on the plasma membrane and mGlyR. (**b**) Representative live-cell fluorescence images of HEK293T cells transfected with mGlyR, without (top) and with (bottom) antimycin A (AA) treatment. mGlyR was labelled using a fluorescent anti-ALFA tag nanobody (GFP channel, left); oxidative stress was visualised using MitoSOX Red (right). Scale bars, 20 μm. (**c**) Quantification of relative fluorescence intensity of MitoSOX (red) and mGlyR (grey) across the cells labelled in **b**. (**d**) Native mass spectrum of mGlyR purified from AA treated cells. Inset: bar chart representing the fold change in relative monomer abundance from *n* = 4 technical replicates; P value is indicated. (**e**) Low *m*/*z* region of MS^2^ spectrum from nTDMS of mGlyR incubated with a 10-fold molar excess of brain lipids, followed by Fenton chemistry (Fe^2+^ and H_2_O_2_) to generate peroxyl radicals *in vitro* (top) or without Fenton treatment (bottom) (quadrupole selection of 6500 *m*/*z*, 1000 *m*/*z* wide, 120 V HCD activation). Isotopic envelopes corresponding to lipid peaks are seen. Lipids were characterised following an ion-trap MS^3^ experiment (**Supplementary Fig. 2-7**) and lengths are annotated from a database search. (**f**) nTDMS spectrum of mGlyR following IRMPD fragmentation from AA-treated cells (top) and untreated cells (bottom). Assigned fragment ions are labelled and mapped onto the structure of mGlyR (PDB: 7SHE). (**g**) Cartoon schematic of the cAMP assay using the GloSensor luminescence system. (**h**) GloSensor Luminescence (RLU) of an untransfected control (light blue 1) and mGlyR-expressing cells without AA treatment (green 3), and control with AA treatment (dark blue 2) and mGlyR-expressing cells with AA treatment (dark green 4). Forskolin was added after 580 s. (**i**) plot depicting the total luminescence for each condition (log_10_-scaled area under the curve in **h**). Error bars represent the standard error of the mean (n=4 biological replicates). Statistical comparisons were performed using a one-way ANOVA and P-values corrected using Tukey post hoc correction; adjusted P values are provided in **Supplementary Table 5**.

To model oxidative stress, HEK293T cells overexpressing mGlyR were treated with the mitochondrial complex III inhibitor antimycin A (AA). AA disrupts electron transport at complex III of the electron transport chain. This leads to upstream electron accumulation and increased electron leakage to molecular oxygen which thereby elevates mitochondrial superoxide production. To optimise conditions for inducing oxidative stress while maintaining cell viability, cytotoxicity was assessed by flow cytometry. We used Annexin-V staining to detect externalised PS (a marker for early apoptosis) and 7-aminoactinomycin D (7-AAD) to identify dead cells. No significant increase in Annexin-V or 7-AAD positive cells was observed following AA treatment (0–60 µM, 1 h), confirming that the oxidative challenge was sub-cytotoxic under these conditions (**Supplementary Fig. 17**). Increased ROS was confirmed by fluorescence microscopy using MitoSOX, a superoxide indicator ^48^. AA induced a marked increase in superoxide signal, extending to the plasma membrane (**Fig. 4b**,**c**). We then established that MitoSOX fluorescence colocalised with the plasma membrane pool of mGlyR using a fluorescently labelled nanobody targeting an N-terminal ALFA tag on the receptor. Importantly, membrane-localised MitoSOX signal was largely absent under basal conditions.

To determine whether ROS exposure alters mGlyR composition, the receptor was purified from these AA-treated cells and analysed by nMS. The spectrum was centred around ∼6000 *m*/*z*, as observed previously, corresponding to a deconvoluted mass of 192,714 ± 552 Da, indistinguishable from the untreated receptor. A minor population of monomeric species (95,815 ± 366 Da) was also observed (**Fig. 4d**); this population of receptors are likely non-functional. The mass of the monomer is ∼600 Da less than half of that of dimer, indicative of loss of a tightly bound lipid at the dimer interface. Relative to untreated mGlyR where monomer abundance was <11%, (**Fig. 1b, Fig. 4d inset**) ROS exposure increased the monomer fraction to ∼24%, representing a 2.2-fold increase. Together, these observations suggest that labile components, such as bound lipids, contribute to dimer stability.

To directly assess the effect of ROS on mGlyR-lipid interactions, delipidated mGlyR was incubated with the brain lipid extract and subsequently subjected to Fenton chemistry (H_2_O_2_/Fe^2+^) to generate ROS prior to MS analysis. Upon ROS exposure, 18 lipid features were detected; however, peaks previously observed in the lower *m*/*z* region, corresponding to PE-P species, were largely absent (**Fig. 4e**). This indicates that ROS selectively disrupts PE-P association with mGlyR. To assess whether this reflects chemical degradation of PE-P, a PE(P-18:0/18:1) standard was subjected to Fenton chemistry before MS analysis. Peaks corresponding to intact PE(P-18:0/18:1) were markedly depleted upon ROS exposure, accompanied by the appearance of new peaks at lower *m*/*z* (**Supplementary Fig. 18**). These features were assigned chemical formulae consistent with oxidative degradation and structural rearrangement. These observations are in-line with radical-mediated oxidative cleavage of the chemically labile vinyl ether bond ^51,52^. Oxidative degradation of PE-P is therefore sufficient to account for its loss of association with mGlyR. Together with the reduction in dimer stability, these findings suggest that PE-P lipids contribute to receptor dimerisation through interactions at, or proximal to, the dimer interface.

To determine whether the PE-P loss reflects covalent modification of the receptor, the mGlyR produced in the AA treated cells was subjected nTDMS. To induce fragmentation of the protein backbone, infrared multiple photon dissociation (IRMPD) was used for activation ^53,54^. Peptide mapping achieved coverage across the transmembrane helices, where covalent lipid adducts would be expected (**Fig. 4f**). Comparison with the fragmentation pattern of untreated mGlyR revealed no evidence of covalent lipid attachment or oxidative receptor modification. We conclude that ROS-induced exposure of mGlyR leads to PE-P depletion but does not lead to stable covalent modification of the protein sequence in the transmembrane region.

Given no sequence modification, we next sought to assess the functional impact of oxidative stress on mGlyR. We examined mGlyR-mediated cAMP signalling using a luciferase-based reporter assay in which luminescence intensity reflects intracellular cAMP levels (**Fig. 4g**). HEK293T cells co-expressing mGlyR, RGS7 and Gβ5 were treated with AA to generate ROS prior to the assay. During the luminescence recording, cells were stimulated with forskolin (at 580 s) to elevate cAMP via activation of adenylyl cyclase, providing a defined baseline from which G_i/o_-mediated inhibition could be measured ^55^ (**Fig. 4h**). We confirmed that, in the absence of ROS, mGlyR-RGS7-Gβ5 led to reduced cAMP accumulation relative to untransfected controls, reflecting G_i/o_-mediated inhibition of adenylyl cyclase. AA pre-treatment further reduced cAMP levels (**Fig. 4h**). This effect was enhanced in mGlyR-RGS7-Gβ5 expressing cells compared to untransfected controls, mimicking the glycine-induced suppression of cAMP ^28^. Quantification of luminescence signal confirmed significantly greater inhibition of cAMP accumulation in mGlyR-expressing cells (**Fig. 4i, Supplementary Table 5**).

Together, these findings support a model in which receptor-associated PE-P lipids act as redox buffers at the mGlyR surface, contributing to both receptor stability and signalling capacity. Oxidative depletion of these lipids, which are selectively enriched in the resting receptor and reduced upon activation coincides with decreased dimer stability, and altered G_i/o_-mediated signalling. These observations suggest that the lipid microenvironment of mGlyR is sensitive to the cellular redox state. Furthermore, it raises the possibility that oxidative modification of receptor-associated lipids provides a mechanism by which the redox environment can modulate both receptor architecture and function.

## Discussion

We developed REVEAL, an automated, untargeted nTDMS strategy for the direct identification and quantification of lipids non-covalently associated with intact, heterogeneous mammalian membrane protein complexes. By resolving specific lipid interactions against a background of over 1,500 components – without prior enrichment or targeting – REVEAL enables assignment of brain lipid interactions to fully modified GPCRs where extensive proteoform heterogeneity precludes conventional nMS-based analysis. The approach is system-agnostic and will be widely applicable as nMS increasingly moves toward the analysis of endogenous mammalian assemblies ^53,56,57^. As such it provides a general framework for mapping non-covalent protein-lipid interactions from tissue-derived samples and tracking how these interactions are remodelled across different functional protein states. Collectively, our findings demonstrate that GPCR-associated lipids are not static structural components, but dynamic interaction partners remodelled by receptor conformation and chemical environment.

Despite sharing a dimeric class C architecture, mGlyR associates with a substantially broader lipid repertoire than mGluR2. This likely reflects differences in TM organisation and inter-protomer spacing: the more open architecture of mGlyR is able to accommodate a wider range of lipid species, potentially permitting lipid interdigitation between helices. By contrast the more compact mGluR2 arrangement imposes greater geometric constraints on lipid binding from the outset. It is also worth noting that not all identified lipids definitively occupy buried or interfacial sites; species such as PC are likely enriched at the outer leaflet and may instead form part of the annular shell surrounding the receptor periphery. Whether individual lipids occupy buried interfacial cavities, inter-helical spaces, or peripheral annular positions remains to be resolved.

Agonism across class C GPCRs is associated with defined rearrangements of the TM domains ^26,58,59^. Consistent with this, mGluR2 exhibits pronounced remodelling of its lipid-binding profile upon agonist binding, coincident with compaction of the inter-protomer pocket. Similarly, glycine agonism of mGlyR is accompanied by marked reduction in associated lipids and relative enrichment of lipids typically found in the inner-leaflet of the plasma membrane. This is consistent with structural reorganisation and increased selectivity for specific lipid chain lengths and degrees of unsaturation. The intermediate lipid profile observed upon L-alanine binding further supports a coupling between conformational engagement and lipid selectivity.

Across all mGlyR states examined, PE-Ps emerge as persistent interaction partners, consistent with reported phospholipid densities within the TM cavities in recent cryo-EM structures ^3,27^. The conservation of these interactions across receptor states suggests that plasmalogen engagement is relatively insensitive to activation-associated conformational changes. Rather, the selective retention of specific PE-P species upon activation, or partial agonism, suggests that conformational changes restrict lipid accessibility, favouring a geometrically and chemically constrained subset of plasmalogens.

Plasmalogens are abundant in neuronal membranes, where they contribute to membrane organisation and redox homeostasis ^51,52,60^. Their vinyl ether bond renders them particularly susceptible to oxidative damage ^51,52,60^, resulting in their preferential depletion under conditions of elevated ROS. The identification of PE-P species as conserved mGlyR interaction partners therefore provides a mechanistic rationale for examining how redox-driven membrane remodelling influences receptor-lipid coupling. Consistent with this, oxidative stress leads to loss of PE-P association, reduced dimer stability and altered G_i/o_-mediated signalling. Although these observations do not establish direct causality, they support a model in which receptor-associated plasmalogens contribute to both receptor architecture and signalling competence, and raise a possibility that oxidative perturbation of plasmalogen-rich membrane environments may provide a physiologically relevant mechanism for modulating mGlyR function **(Fig. 5)**.

**Figure 5.**
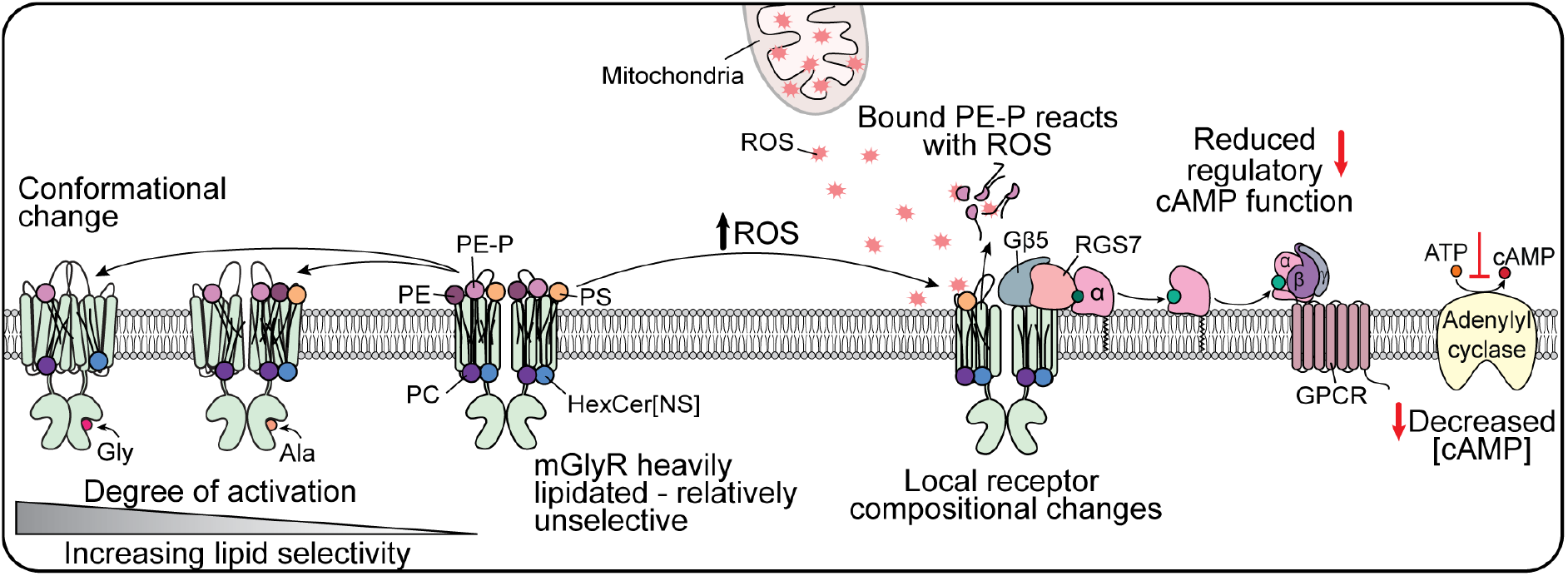
Summary of mGlyR-lipid interactions detected under apo, agonist-bound, and oxidative stress conditions. Under apo conditions, mGlyR binds five lipid classes spanning a range of acyl chain lengths and degrees of unsaturation. Upon binding of L-alanine or the agonist glycine, mGlyR exhibits increased lipid selectivity. The effect is more pronounced for glycine, consistent with its established role in eliciting downstream signalling. Under oxidative stress conditions, PE-P lipids normally associated with mGlyR are no longer detected, due to radical-mediated oxidation and degradation. Concomitantly, mGlyR exhibits localised structural changes and a reduced cAMP response.

More broadly, these findings position membrane lipid composition, and its redox sensitivity, as a regulatory axis for GPCR signalling that is distinct from, yet coupled to, ligand binding and conformational control. REVEAL provides the methodological foundation to interrogate this axis systematically across the membrane proteome. It also enables us to determine how lipid environments are reshaped by the metabolic and oxidative conditions of pathological states. Together, this work establishes a link between lipid chemistry, receptor architecture and class C GPCR signalling.

## Acknowledgements

The authors thank J. L. Bennett, D. Wu, S. Song and M. R. L. Williams for useful discussions. The authors are grateful to colleagues at Thermo Fisher Scientific Life Sciences Mass Spectrometry division for providing critical support and infrastructure to facilitate the Orbitrap Ascend instrument modifications. Vlad Zabrouskov (Thermo Fisher Scientific, San Jose, CA) and Alexander Makarov (Thermo Fisher Scientific Bremen, Germany) is thanked for useful discussions. We thank R. Hedley and V. Tsioligka for providing technical assistance with fluorescence-assisted cell sorting at The Don Mason Facility of Flow Cytometry, Sir William Dunn School of Pathology, University of Oxford. The authors gratefully acknowledge the Micron Advanced Bioimaging Facility, University of Oxford (supported by Wellcome Strategic Awards 091911/B/10/Z and 107457/Z/15/Z) for their support & assistance in this work.

## Data availability

All mass spectrometry data have been deposited to the ProteomeXchange Consortium via the PRIDE ^61^ partner repository with the dataset identifier PXD077927. Plasmids and cell lines generated in this study will be made available upon reasonable request to the corresponding authors (Tarick El-Baba or Carol Robinson).

## Materials and methods

### Reagents

n-dodecyl β-D-maltoside (DDM) and n-decyl-β-D-maltoside (DM) were purchased from Anatrace. Brain polar lipid extract was purchased from Avanti Research. All other chemicals were purchased from Merck Millipore, unless specified.

### Cell culture

HEK293S GnTI^-/-^ (ATCC-CRL-3022) were obtained from LGC standards. Cells were cultured as adherent monolayers in 1X Dulbecco’s modified eagle medium:Ham’s F12 (DMEM:F12) (Life Technologies, Thermo Fisher Scientific) supplemented with 10% fetal bovine serum (FBS) (Life Technologies, Thermo Fisher Scientific), and 1X non-essential amino acids (Life Technologies, Thermo Fisher Scientific).

### Expression and purification of mGluR2 and mGlyR

Synthetic, codon optimised DNA fragments encoding mGluR2 (residues 1 – 872) and mGlyR (residues 1–775) were purchased from Integrated DNA Technologies (IDT). Both constructs contained C-terminal tags designed for affinity purification; the mGluR2 construct included an AVI and FLAG tag (ENLYFQGSGGSGGSGLNDIFEAQKIEWHEGGSGGSDYKDDDDK) and the mGlyR construct had Twin-Strep and FLAG tags. (ENLYFQGSWSHPQFEKGGGSGGGSGGGSWSHPQFEKGGGSGGGSGGGSDYK DDDDK). The DNA fragments were cloned into a lentiviral transfer plasmid (a kind gift from J. Elegheert and R. Aricescu, Addgene plasmid #113884) and lentiviruses were generated as described ^62^. 3 days after infection, the supernatant (18 mL) containing high-titre lentiviral particles was used to infect HEK293 GNTI^-/-^ cells as described ^62-64^. Cells positive for eGFP were separated from non-transduced cells using fluorescence-activated cell sorting (FACS) at the William Dunn School of Pathology, University of Oxford. Polyclonal cell lines were adapted to suspension using 1X FreeStyle (Thermo Fisher Scientific) with 1% FBS (v/v). When a density of ∼3 × 10^6^ cells mL^-1^ was reached, cells were induced with tetracycline (1 µg mL^-1^). Protein expression was boosted with sodium butyrate (5 mM) after 24 hr. Cells were harvested ∼48 hr after induction by centrifugation (1,000 ×g, 20 min, 4 °C), snap frozen in LN2, and stored at -80 °C.

Cells were thawed on ice, resuspended in HNG buffer (50 mM HEPES pH 8.0, 150 mM NaCl, 5% glycerol) supplemented with EDTA-free protease inhibitor tablets (Roche), and lysed using a microfluidizer. Debris was removed by centrifugation (14,000 ×g, 20 min, 4 °C) and the membrane fraction was collected by ultracentrifugation of the supernatant (100,000 ×g, 1 h, 4 °C). The pelleted membrane was then resuspended in HNG, snap frozen in LN2, and stored at -80 °C until use.

The membranes were solubilized with HNG buffer supplemented with 1% (w/v) DDM/0.1% cholesteryl hemi-succinate (CHS) for 2 hr at 4°C with rotation. The solubilized membranes were clarified by centrifugation (14,000 ×g, 20 min, 4 °C) and the supernatant was incubated with anti-DYKDDDK magnetic agarose resin (Life Technologies, Thermo Fisher Scientific) for 2 hr. The lysate was separated from the magnetic particles with a magnet. The immobilized proteins were then washed with 20 resin volumes of HNG buffer supplemented with 0.017%/0.0017% DDM/CHS prior to elution using 1 mg mL^-1^ synthetic DYKDDDDK peptide (Genscript). mGluR2 and mGlyR were concentrated to ∼10 µM (dimer concentration) using a 100 kDa MWCO concentrator, aliquoted, snap frozen in LN2, and stored at -80 °C until use.

For delipidating purifications, the membranes were solubilized with HNG buffer supplemented with 2% (w/v) DDM/0.2% CHS for 2 hr at 4°C with rotation. The solubilized membranes were clarified by centrifugation (14,000 ×g, 20 min, 4 °C) and the supernatant was incubated with anti-DYKDDDK magnetic agarose resin (Life Technologies, Thermo Fisher Scientific) for 1 hr. The lysate was separated from the magnetic particles with a magnet. To delipidate, the immobilised proteins were incubated with HNG buffer supplemented with 2% DDM/0.2% CHS for 12 hr with rotation at 4 °C. The delipidated proteins were then washed with 20 resin volumes of HNG buffer supplemented with 0.017%/0.0017% DDM/CHS prior to elution using 1 mg mL^-1^ synthetic DYKDDDDK peptide (Genscript). mGluR2 and mGlyR were was concentrated to ∼10 µM (dimer concentration) using a 100 kDa MWCO concentrator, aliquoted, snap frozen in LN2, and stored at -80 °C until use.

For antimycin A treated purifications, cells were induced with tetracycline (1 µg mL^-1^) when a density of ∼3 × 10^6^ cells mL^-1^ was reached. Protein expression was boosted with sodium butyrate (5 mM) after 24 hr. Antimycin A (18 µM) was added 1 hr before harvesting. Cells were harvested, and purified as above.

### Protein-lipid binding

For mGluR2 and mGlyR-brain lipid binding experiments, delipidated mGluR2/mGlyR (5 µM dimer) were combined with a 10-fold molar excess of Brain polar lipid extract dissolved in 400 mM ammonium acetate (pH 7.4) supplemented with 0.17%/0.017% DM/CHS. The protein-lipid mixture was incubated at room temperature for 30 min prior to analysis.

### Native mass spectrometry (nMS)

A modified Orbitrap Tribrid Ascend mass spectrometer (Thermo Fisher Scientific) equipped with a high mass quadrupole mass filter and infrared laser directed into the linear ion trap, a design described previously ^53^, was used to acquire native mass spectra of mGluR2 and mGlyR. mGlyR and mGluR2 (5 μM dimer) were buffer exchanged into 400 mM ammonium acetate supplemented with 0.17%/0.017% DM/CHS twice using Zeba desalting column (Pierce) (75 µL, 40 kDa MWCO). ∼2-3 µL of buffer exchanged protein was loaded into a gold-coated nanoelectrospray capillary prepared in house ^65^. All spectra were collected in positive ion mode. Typical electrospray ionisation voltages were held between ∼0.9-1.2 kV relative to the instrument orifice (heated to ∼100-200 °C). The source collision energy was held at 250 V (with a source compensation of 0.01 – 0.05%) to facilitate ejection of the membrane proteins from the proteomicelle. The instrument was operated in high pressure mode (20 mTorr in both ion routing multipoles). A resolution setting of 30,000 (at *m*/*z* 200) was used. To generate native mass spectra for mGluR2 and mGlyR, a stage of infrared micelle dissociation, described previously ^53,54,57^, was used. Typical irradiation times were 5 ms with 6 W output power of the 60 W CO_2_ laser.

### nTDMS

nTDMS experiments were carried out as previously described ^66^ using the same modified Orbitrap Tribrid Ascend mass spectrometer (Thermo Fisher Scientific).

For lipid release MS^2^ experiments, precursors were manually selected using the high *m*/*z* quadrupole using a 100 *m*/*z* window for mGlyR and mGluR2. The selected ions were activated with 100-120 V absolute higher-energy collisional dissociation (HCD) energy in the front HCD cell to promote dissociation of lipids. The dissociated lipid ions were detected in the Orbitrap at a resolution of 30,000 (at *m*/*z* 200). Spectra represent the average of 100–999 individual 1 s acquisitions (automated gain control = 1000%).

The MS^3^ identification of lipid species used the targeted MS^2^ spectra as precursor species. Individual lipid peaks were manually isolated in the ion trap (2 *m*/*z* isolation window) prior to 10 ms resonance collision induced dissociation (CID) at 20-30%. Resulting spectra were recorded in the ion trap in positive and/or negative mode (Turbo or Normal scan rate).

For fragmentation of mGlyR, ions within an *m*/*z* range of interest were isolated in the ion trap using a selection window of 100 *m*/*z* wide and a q value of 0.1. Isolated populations were subjected to infrared multiple photon dissociation for 10 ms using 12% of the maximum laser output (7.2 W) to induce fragmentation. Fragment ions were detected in the Orbitrap mass analyser using a resolving power of 240,000 (at *m*/*z* 200). Maximum injection times were set to 1 s (with a 100% AGC target) and averaged for a maximum number of individual scans (999 scans). Fragments were manually assigned using the targeted top-down module (TDValidator) in ProSight Native v1.0.25339 (Proteinaceous) ^67^ using the sequence of the expressed receptor with the signal peptide removed as the input. Spectra were annotated using a signal-to-noise cut-off of 3.0, a maximum ppm tolerance of 10 ppm, a sub-ppm tolerance of 5 ppm, a cluster tolerance of 0.35 and a minimum score of 0.6 using the distribution generator Mercury7.

### REVEAL

The REVEAL experiments were carried out on the modified Orbitrap Tribrid Ascend mass spectrometer. The automated method was written and executed through XCalibur (version 4.6.67.17). An example method is provided in the **Supplementary Information (Supplementary Fig. 19)**. Prior to executing the method, a precursor (MS^1^) spectrum was recorded for ions between *m*/*z* 2000 to 8000. Upon method execution, regions of the MS^1^ spectrum were sequentially selected in 10 *m*/*z* non-overlapping increments using the quadrupole, and all ions falling within the selection window were subjected to dissociation using HCD (100-120 V). MS^2^ spectra were recorded between *m*/*z* 685–920 for each window. Precursors were between *m*/*z* 5000-8000 for mGluR2 and *m*/*z* 4000–8000 for mGlyR. MS^1^ and MS^2^ spectra were recorded at a resolution of 30,000 (at *m*/*z* 200). Typical instrument parameters include electrospray ionisation voltages held between 1.1-1.3 kV relative to the instrument orifice (heated to ∼100-200 °C); 250 V source collision energy, source CID compensation scaling factor 0.01-0.02% to facilitate ejection of the membrane protein from the proteomicelle; 1000 ms injection time and 100% AGC target. The instrument was operated in high pressure mode (20 mtorr in both ion routing multipoles).

### Lipidomic analysis of the brain polar lipids

Lipidomics analysis of a brain polar lipid extract was carried out as previously described ^3^. Briefly, dried lipids were resuspended in 80% mobile phase A (acetonitrile/water, 60:40 [vol/vol] containing 10 mM ammonium formate and 0.1% [vol/vol] formic acid) and 20% mobile phase B (isopropanol/acetonitrile, 90:10, [vol/vol] containing 10 mM ammonium formate and 0.1% [vol/vol] formic acid). Lipids were separated by liquid chromatography on a C18 column (Acclaim PepMap 100, C18, 75 µM × 15cm, Thermo Scientific) using a Dionex UltiMate 3000 RSLC system and analysed by tandem mass spectrometry (LC-MS/MS). Eluted lipids were analysed on Orbitrap Eclipse mass spectrometer operated in data-dependent acquisition mode in both positive and negative modes. Raw data was processed and lipid species identified using LipiDex ^68^ and distinct chain lengths were summed for each class. Plasmalogen species were quantified using MZmine3 ^69^, and their intensities were expressed relative to total plasmalogen signal.

### Expression and purification of ALFA nanobody

An anti-ALFA nanobody (NbALFA-CE variant) was cloned into a pET-15b vector and expressed in *E. coli* BL21(DE3). Following transformation, cells were plated on LB agar supplemented with 100 µg mL^-1^ ampicillin and grown overnight. Single colonies were used to inoculate a 100 mL starter culture in LB containing ampicillin (100 µg mL^-1^) and 1% glucose, which was subsequently expanded into 1 L of the same medium. Cultures were grown to OD_600_ = 0.4-0.6, induced with 1 mM IPTG, and incubated overnight at 28°C.

Cells were harvested by centrifugation (5,000 × g, 4°C), resuspended in TES buffer (20 mM Tris, pH 8.0 at 4°C, 500 µM EDTA, 500 mM sucrose), and incubated for 1 h at 4°C with rotation. Periplasmic extraction was performed by osmotic shock through addition of an equal volume of 0.25× TES buffer, followed by incubation for 1 h at 4°C. The lysate was clarified by centrifugation (20,000 × g, 1 h, 4°C).

The periplasmic fraction was applied to Ni-NTA resin pre-equilibrated in HNG buffer (50 mM HEPES, pH 8.0, 150 mM NaCl, 5% glycerol). After washing with HNG supplemented with 20 mM imidazole, bound nanobody was eluted with HNG containing 300 mM imidazole. Eluted fractions were desalted using a PD-10 column (Cytiva) according to the manufacturer’s instructions and concentrated to 2 mg mL^-1^.

For fluorescent labelling, nanobodies were reacted with NHS-fluorescein (Thermo Fisher Scientific) following the manufacturer’s protocol. Briefly, NHS-fluorescein was dissolved in DMSO and added to the protein solution while maintaining the final DMSO concentration below 1%. The reaction was incubated for 1 h at room temperature, and excess dye was removed using a Pierce desalting column (4 kDa MWCO). The degree of labelling was assessed by denaturing mass spectrometry. Labelled nanobodies were aliquoted, flash-frozen in liquid nitrogen, and stored at -80°C.

### Live cell fluorescence microscopy

HEK293T cells were transiently transfected with a pcDNA3.1 construct encoding mGlyR containing an N-terminal ALFA tag using Lipofectamine 2000, according to the manufacturer’s instructions. At 24 h post-transfection, cells were treated with 18 µM antimycin A for 1 hr at 37 °C, washed with PBS, and then incubated with 1 µM MitoSOX Red (Invitrogen, M36008) and fluorescent ALFA nanobody for 30 min at 37 °C.

Prior to imaging, the culture medium was replaced with FluoroBrite DMEM (Life Technologies) supplemented with 20 mM HEPES (pH 7.4). Fluorescence imaging was performed on a Leica Thunder inverted microscope using either a 40× objective or a 100× oil-immersion objective. Cells were maintained at 37 °C in a humidified atmosphere with 5% CO_2_ during acquisition. Images were processed using Fiji (version 1.54p).

### Flow cytometry

HEK293T cells were seeded in a 24-well plate and treated with increasing concentrations of antimycin A (0–60 µM) for 1 h at 37 °C. Following treatment, both floating and adherent cells were collected by centrifugation (300 ×g, 5 min). Cells were resuspended in Annexin V binding buffer (10 mM HEPES pH 7.4, 140 mM NaCl, 2.5mM CaCl_2_).

Cells were stained with Annexin V-488 (Alexa Fluor 488) and 7-aminoactinomycin D (7-AAD) according to the manufacturer’s instructions and incubated for 15 min at room temperature in the dark. Following incubation, samples were analysed by flow cytometry.

Data was acquired using a BD LSR Fortessa X-20 flow cytometer (BD Biosciences). Forward and side scatter were used to gate live cells and exclude debris. 10,000 events on P2 (single cells) were collected per sample. Data were analysed using FlowJo software (version 10).

### Fenton chemistry

PE(P-18:0/18:1) was purchased from Avanti Research. PE(P-18:0/18:1) (5 µM in MeOH:H_2_O) was subjected to Fenton oxidation by adding iron(II) sulfate (10 µM stock in H_2_O) and hydrogen peroxide (100 µM stock in H_2_O) in a 3:1:1 (v/v/v) ratio (lipid:Fe^2+^:H_2_O_2_) ^70^. The reaction was incubated at room temperature (10 min) before nMS analysis in positive mode on an Orbitrap Eclipse Tribrid mass spectrometer (Thermo Fisher Scientific). MS^2^ experiments were carried out to identify lipids, as above: individual peaks were manually isolated in the ion trap (2 *m*/*z* isolation window) prior to 10 ms resonance collision induced dissociation (CID) at 20-30%. Resulting spectra were recorded in the ion trap in positive mode.

### In vitro Fenton chemistry

For mGlyR-brain lipid oxidation experiments, delipidated mGlyR (5 µM dimer) were combined with a 10-fold molar excess of brain polar lipid extract dissolved in 400 mM ammonium acetate (pH 7.4) supplemented with 0.17%/0.017% DM/CHS. The protein-lipid mixture was incubated at room temperature for 30 min. The mixture was then subjected to Fenton oxidation by adding iron(II) sulfate (10 µM stock in H_2_O) and hydrogen peroxide (100 µM stock in H_2_O) in a 3:1:1 volume ratio (protein/lipid mixture:Fe^2+^:H_2_O_2_). The reaction was incubated at room temperature for 10 minutes before nMS analysis.

### cAMP accumulation assay

pcDNA3.1 plasmids encoding mGlyR, RGS7, Gβ5, and the luciferase-based cAMP biosensor pGloSensor-22F (Promega) were mixed at a 1:1:1:1 ratio and transiently transfected into HEK293T cells using Lipofectamine 2000 according to the manufacturer’s protocol. 24 hr after transfection, cells were treated with 18 µM antimycin A (ROS conditions) or DMSO alone (vehicle control) for 1 hr at 37 °C, with final DMSO concentration kept constant (<1 % v/v). Culture media was then replaced with CO_2_-independent medium supplemented with 10% FBS and 2% (v/v) Glosensor assay reagent (Promega) as per the manufacturers protocol, and the plate was incubated in the dark for 1hr at room temperature.

Luminescence was measured using a FLUOstar Omega Microplate Reader (BMG Labtech). After 10 measurement cycles (580 s), forskolin was added to a final concentration of 100 µM.

### Data analysis

Native mass spectra and lipid ion chromatograms were extracted from QualBrowser (Xcalibur version 4.5.474.0) and imported into OriginPro 2024b (version 10.1.5.132). MS^1^ spectra were normalized between 0 and 1 for plotting. Extracted ion chromatograms were manually converted from the time domain into *m*/*z* to reconstruct the “precursor ion spectra”. Relative quantification was carried out by integrating areas of interest using the peak integrator tool in OriginPro. For lipid class identification, the manually assigned MS^3^ fragmentation of the lipids were searched against the LIPID MAPS database ^71^. Protein electrostatics were measured in ChimeraX ^72^. TM helices were measured from the top and bottom of each helix. Measurements were taken from C-α using ChimeraX ^72^. Protein interfaces and associated free energies were analysed using the PDBePISA server ^73^. Protein surface pockets and cavities were identified using CASTpFold (Computed Atlas of Surface Topography of Proteins for Folded Structures), which calculates solvent-accessible areas and volumes based on the 3D structure ^74^. Default parameters were applied.

