## Supplemental Info for "Defining state-selective lipid binding to brain GPCRs - introducing REVEAL"

**Supplementary Table 1. Expected molecular weights of recombinant proteins used in this study.** Expected masses only consider N-glycans (1216 Da) and signal peptide cleavage. Masses reported as average ± s.d. deconvoluted from ≥3 of the most abundant charge state peaks (*viz.* most abundant proteoform).

| | **Sequence mass[1]** | **Expected mass[1,2]** | **Measured mass** |
|---|---|---|---|
| mGlyR | 179,375 Da | 191,535 Da | 192,075 ± 161 Da |
| mGluR2 | 196,620 Da | 208,780 Da | 210,808 ± 206 Da |

[1] Assumes full cleavage of signal peptide
[2] Includes ten $Man_5GlcNAc_2$ glycans (five per monomer) for both proteins

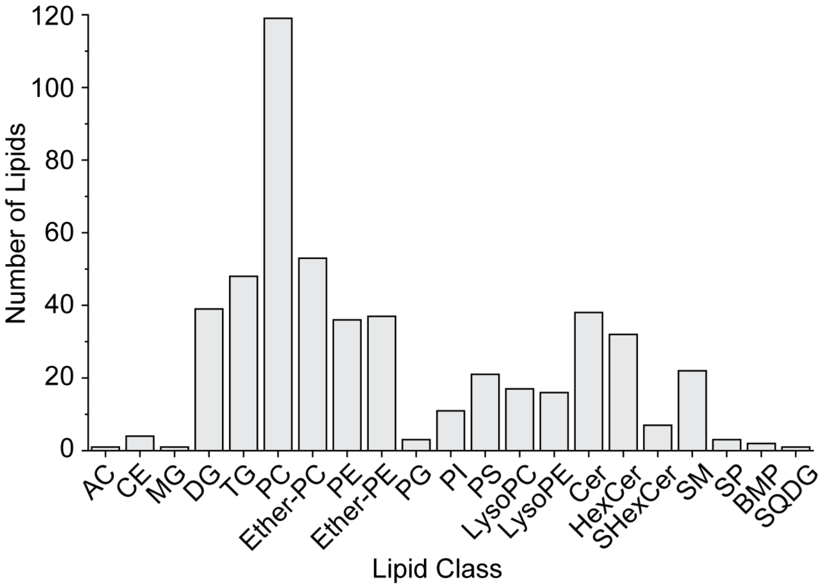

**Supplementary Figure 1. Lipidomic analysis of brain lipid polar extract.**
Bar graph showing the number of unique lipid species detected in each lipid class. Lipid species are defined by distinct fatty acid compositions. Abbreviations: AC, acylcarnitine; CE, cholesteryl ester; MG, monoacylglycerol; DG, diacylglycerol; TG, triacylglycerol; PC, phosphatidylcholine; ether-PC, ether-linked phosphatidylcholine; PE, phosphatidylethanolamine; ether-PE, ether-linked phosphatidylethanolamine; PG, phosphatidylglycerol; PI, phosphatidylinositol; PS, phosphatidylserine; lysoPC, lysophosphatidylcholine; lysoPE, lysophosphatidylethanolamine; Cer, ceramide; HexCer, hexosylceramide; SHexCer, sulfohexosylceramide; SM, sphingomyelin; SP, sphingoid base; BMP, bis(monoacylglycero)phosphate; SQDG, sulfoquinovosyldiacylglycerol.

**Supplementary Table 2. Assigned peak ID of each released lipid identified in this study.**
Identities were made using $MS^3$.

| *m/z* | Peak Identity |
|---|---|
| **700.53** | PE(P-34:2) $[M+H]^+$ |
| **702.54** | PE(P-34:1) $[M+H]^+$ |
| **716.52** | PE(P-35:1) $[M+H]^+$ |
| **718.54** | PE(34:1) and PC(31:1) $[M+H]^+$ |
| **724.53** | PE(P-36:4) $[M+H]^+$ |
| **726.54** | PE(P-36:3) $[M+H]^+$ |
| **728.56** | PE(P-36:2) $[M+H]^+$ |
| **730.57** | PE(P-36:1) $[M+H]^+$ |
| **732.56** | PE(P-36:0) $[M+H]^+$ |
| **740.52** | PE(36:4) $[M+H]^+$ |
| **742.53** | PE(P-37:2) $[M+H]^+$ |
| **744.55** | PE(36:2) $[M+H]^+$ |
| **746.57** | PE(36:1) $[M+H]^+$ |
| **750.54** | PE(P-38:5) $[M+H]^+$ and PE(35:2) $[M+Na]^+$ |
| **752.56** | PE(P-38:4) $[M+H]^+$ and PE(35:1) $[M+Na]^+$ |
| **756.59** | PE(P-38:2) $[M+H]^+$ |
| **758.61** | PE(P-38:1) and PC(34:2) $[M+H]^+$ |
| **760.59** | PC(34:1) $[M+H]^+$ |
| **764.52** | HexCer[NS](d18:1_21:1) $[M+H]^+$ and PE(36:3) $[M+Na]^+$ |
| **766.54** | PE(36:2) $[M+Na]^+$ |
| **768.55** | PE(36:1) $[M+Na]^+$ |
| **772.59** | PE(38:2) and PC(35:2) $[M+H]^+$ |
| **774.53** | PE(P-40:7) and PC(35:1) $[M+H]^+$ |
| **776.56** | PE(P-40:6) and PC(35:0) $[M+H]^+$ |
| **778.57** | PE(P-40:5) and PC(36:6) $[M+H]^+$ |
| **780.59** | HexCer[NS](d18:1_22:2) and PE(P-40:4) $[M+H]^+$ and PC(34:2) and PE(37:2) $[M+Na]^+$ |
| **782.61** | PE(P-40:3) $[M+H]^+$ |
| **784.62** | PE(P-40:2) and PC(36:3) $[M+H]^+$ |
| **786.63** | PC(36:2) $[M+H]^+$ |
| **788.54** | PC(36:1) $[M+H]^+$ |
| **790.56** | PS(36:1) $[M+H]^+$ |
| **792.56** | HexCer[NS](d18:1_23:3) $[M+H]^+$ and PE(38:3) $[M+Na]^+$ |
| **794.57** | HexCer[NS](d18:1_23:2) $[M+H]^+$ and PE(38:2) $[M+Na]^+$ |
| **796.58** | HexCer[NS](d18:1_23:1) $[M+H]^+$ and PE(38:1) $[M+Na]^+$ |
| **808.67** | HexCer[NS](d18:1_24:2) $[M+H]^+$ |
| **810.50** | PS(36:2) $[M+Na]^+$ |
| **810.60** | HexCer[NS](d18:1_24:2) $[M+H]^+$ |
| **812.54** | PC(36:0) $[M+Na]^+$ |
| **814.52** | PE(40:6) and PC(37:6) $[M+Na]^+$ |
| **816.57** | HexCer[NS](d18:1_25:5) and PC(38:1) $[M+H]^+$ and PE(40:5)$[M+Na]^+$ |
| **820.63** | HexCer[NS](d18:1_25:3) $[M+H]^+$ |
| **822.64** | HexCer[NS](d18:1_25:2) $[M+H]^+$ |
| **832.67** | PC(40:7) $[M+H]^+$ |
| **834.68** | PC(40:6) $[M+H]^+$ |
| **836.54** | HexCer[NS](d14:0_29:3) $[M+H]^+$ and PE(40:5) and PC(38:2) $[M+Na]^+$ |
| **838.54** | HexCer[NS](d14:0_29:2) $[M+H]^+$ |
| **840.54** | HexCer[NS](d14:0_29:1) $[M+H]^+$ |
| **848.66** | PC(41:6) $[M+H]^+$ |
| **850.68** | PC(41:5) $[M+H]^+$ |
| **909.55** | PI(39:0) $[M+H]^+$ |

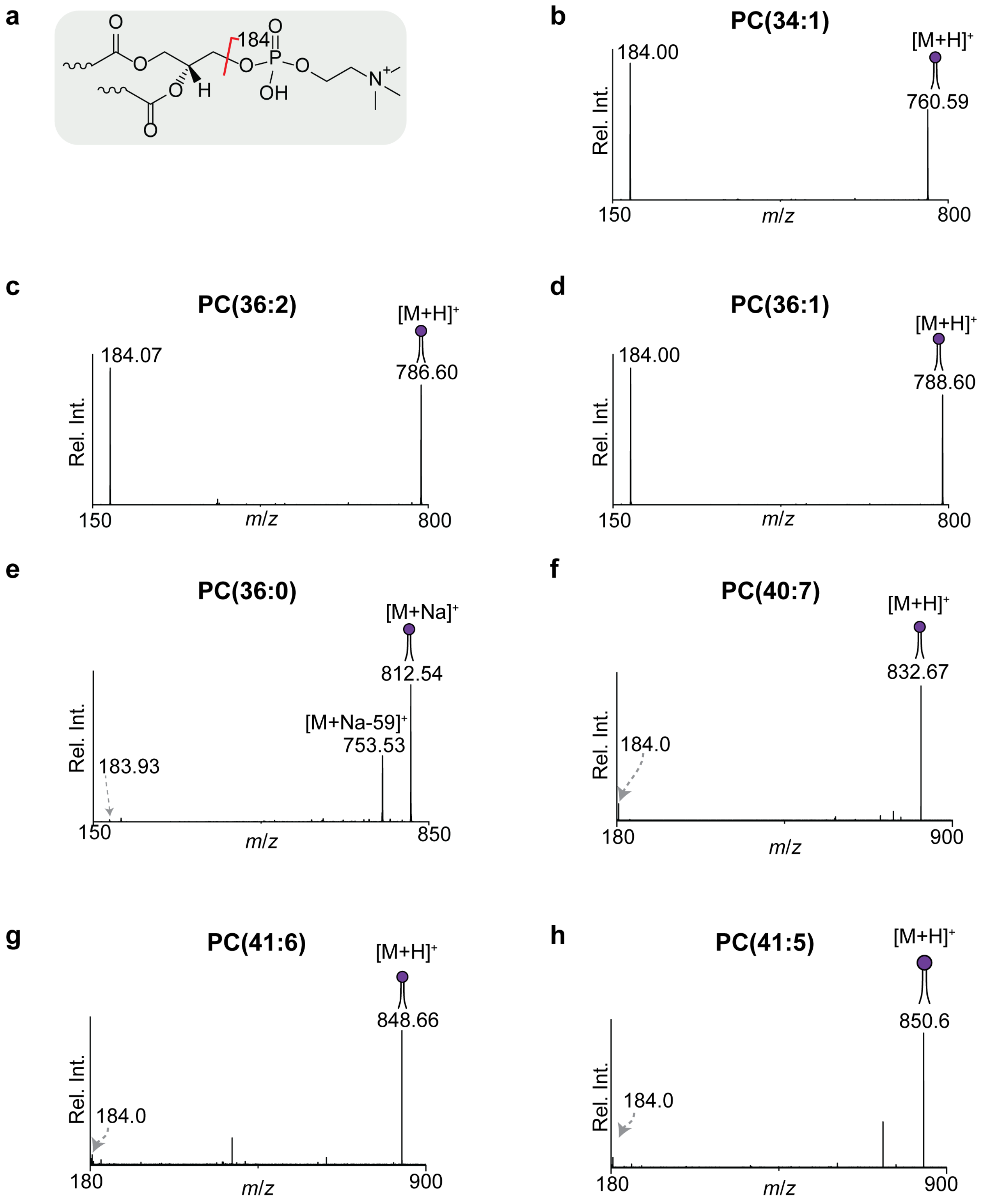

**Supplementary Figure 2. Assignment of PC lipids.**

**a,** Common fragmentation pathway for PC in positive mode, to show characteristic fragments. Headgroup loss from protonated PC results in a peak at *m/z* 184.0. **b-h,** Ion-trap CID fragmentation of PC species identified in mGluR2/mGlyR. Spectra were collected as MS$^3$ from mGluR2/mGlyR incubated with brain lipids.

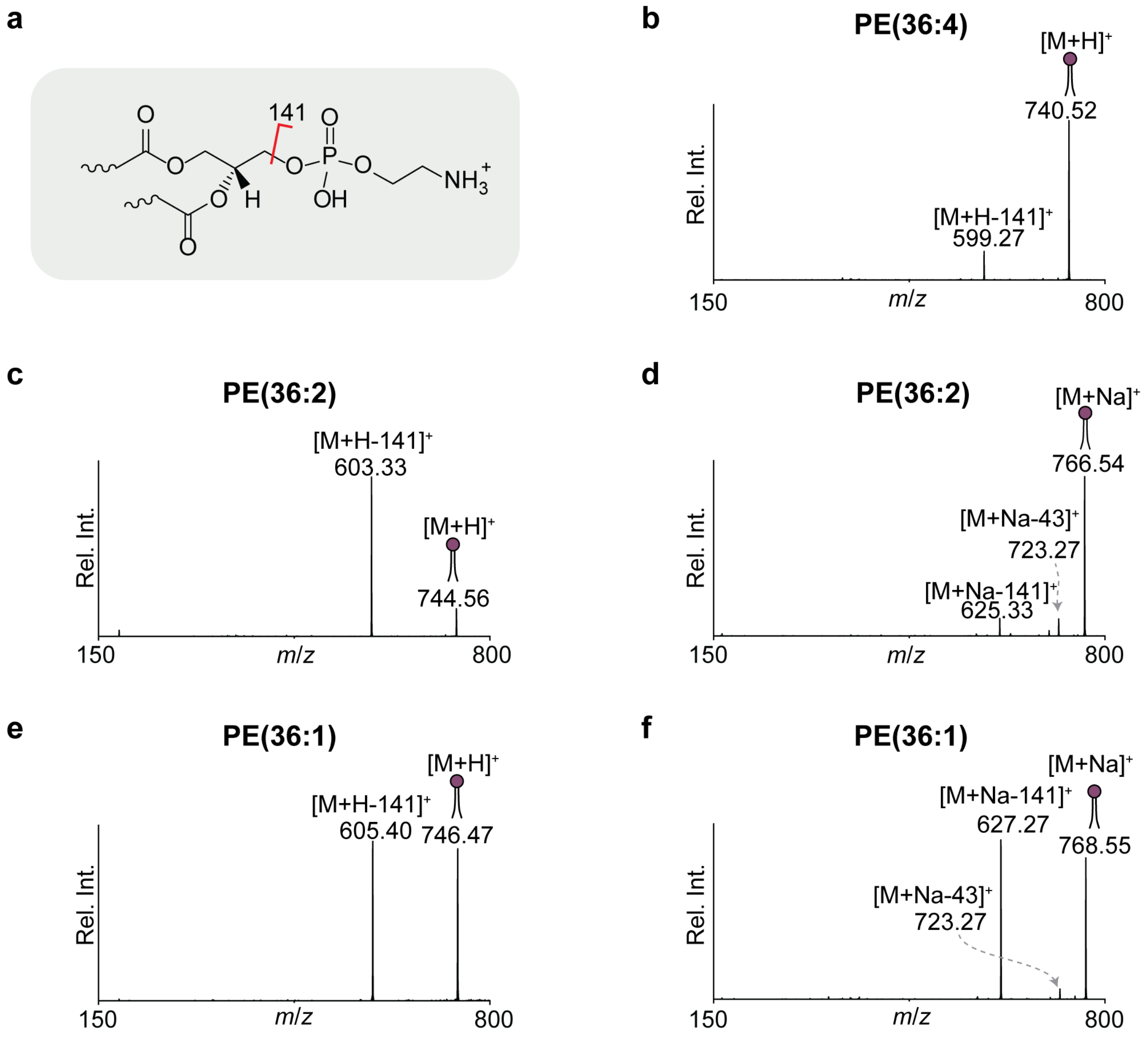

**Supplementary Figure 3. Assignment of PE lipids.**
**a,** Common fragmentation pathway for PE in positive mode, to show characteristic fragments. Headgroup loss from protonated PE results in a peak at 141.0 Da less than the precursor. **b-f,** Ion-trap CID fragmentation of PE species identified in mGluR2/mGlyR. Spectra were collected as $MS^3$ from mGluR2/mGlyR incubated with brain lipids.

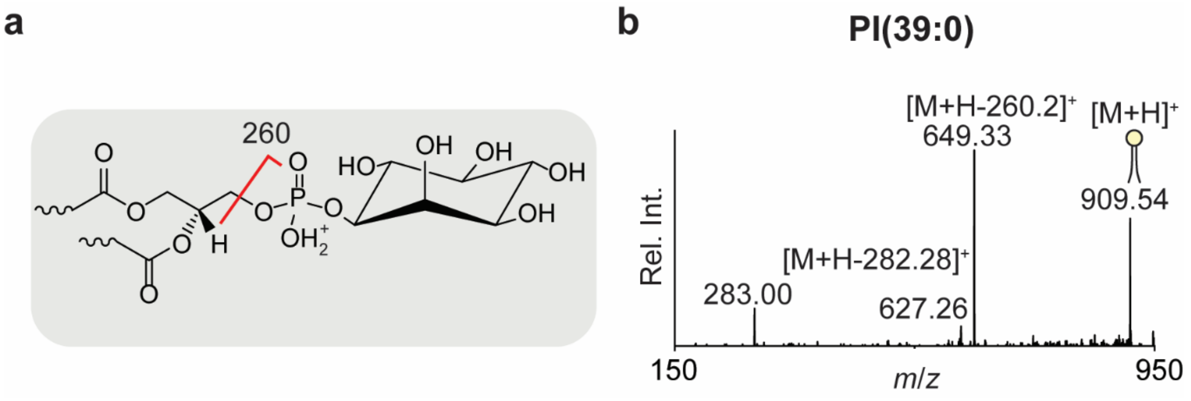

**Supplementary Figure 4. Assignment of PI lipid.**
**a,** Common fragmentation pathway for PI in positive mode, to show characteristic fragment. Headgroup loss from protonated PI results in a peak at *m*/*z* 260.0. **b,** Ion-trap CID fragmentation of PI species identified in mGlyR. Spectra were collected as $MS^3$ from mGlyR incubated with brain lipids.

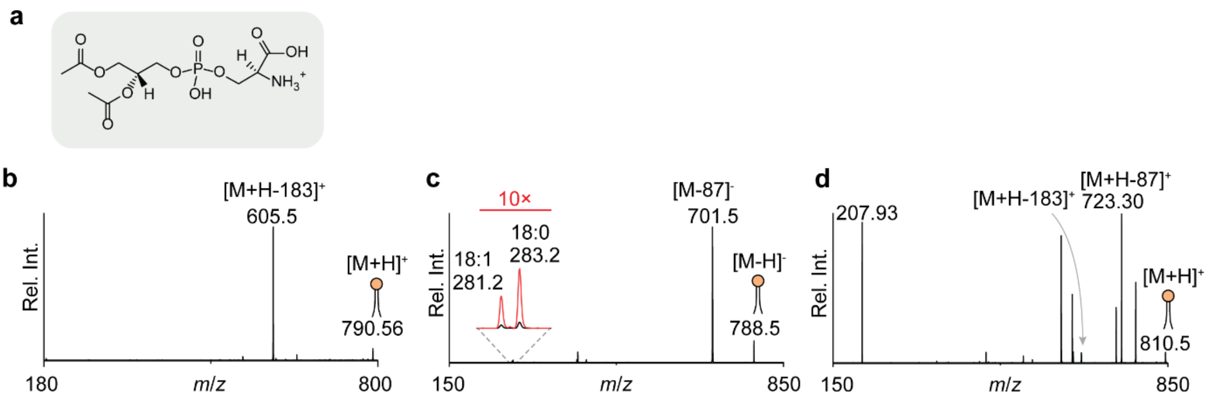

**Supplementary Figure 5. Assignment of PS lipids.**
**a,** Structure of PS headgroup. **b**, Positive mode ion-trap CID fragmentation of PS(36:1). **c**, Negative mode ion-trap CID fragmentation of PS(36:1). A peak 87 Da less than the precursor indicates a serine loss, and peaks for the fatty acid tails are also observed. **d**, Positive mode ion-trap CID fragmentation of PS(36:2). Positive mode spectra were collected as $MS^3$ from mGluR2/mGlyR incubated with brain lipids, negative mode spectrum was collected as an $MS^2$ directly from the brain lipid extract.

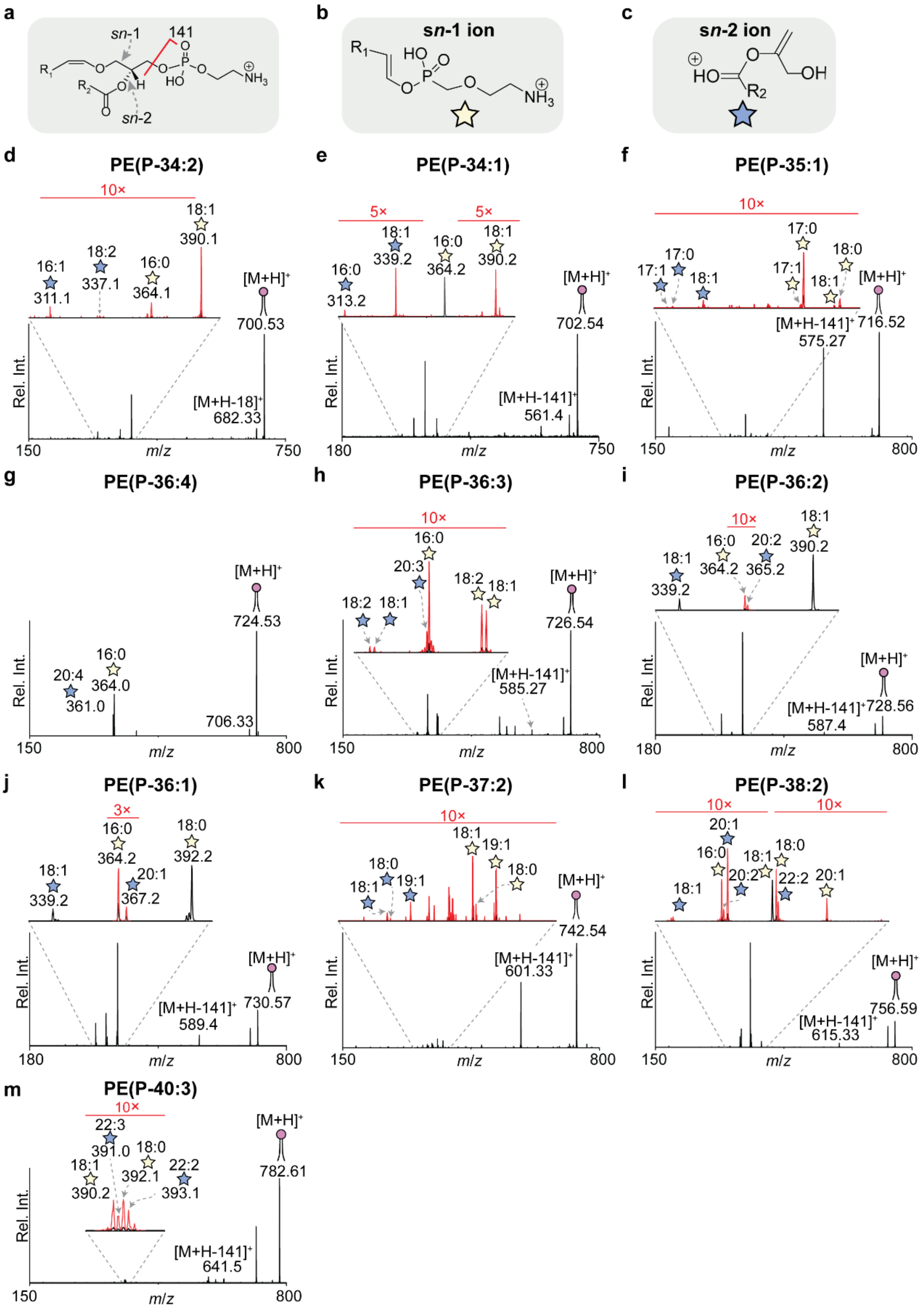

**Supplementary Figure 6. Assignment of plasmenyl-PE lipids.**
**a,** Structure of plasmenyl-PE showing vinyl ether linkage at the *sn*-1 position and the characteristic fragment corresponding to head group loss. Neutral headgroup loss from protonated PE results in a peak at -141 Da. **b,** Literature mechanism for the rearrangement of plasmenyl-PE to form product ions characteristic of the *sn*-2 position (blue stars) (*1-4*). **c,** Literature mechanism for the rearrangement of plasmenyl-PE to form characteristic product ions of the sn-1 position (yellow stars) (*1-4*). **d-m,** Ion-trap CID fragmentation spectra in positive mode for plasmenyl-PE species identified in mGluR2/mGlyR. Spectra were collected as $MS^3$ from mGluR2/mGlyR incubated with brain lipids.

**Supplementary Table 3. Expected *m*/*z* for s*n*-1 and s*n*-2 products of different plasmenyl-PE fatty acid lengths.**
Used to aid assignment of **Supplementary Fig. 6**.

| **Chain Length** | **s*n*-1 (*m*/*z*)** | **s*n*-2 (*m*/*z*)** |
|---|---|---|
| 14:0 | 336.2 | 285.2 |
| 16:0 | 364.2 | 313.2 |
| 17:0 | 378.2 | 327.2 |
| 18:0 | 392.2 | 341.2 |
| 18:1 | 390.2 | 339.2 |
| 18:2 | 388.2 | 337.2 |
| 19:0 | 406.2 | 355.2 |
| 19:1 | 404.2 | 353.2 |
| 20:0 | 420.2 | 369.2 |
| 20:1 | 418.2 | 367.2 |
| 20:2 | 416.2 | 365.2 |
| 20:3 | 414.2 | 363.2 |
| 20:4 | 412.2 | 361.2 |
| 21:0 | 434.2 | 383.2 |
| 21:1 | 432.2 | 381.2 |
| 21:2 | 430.2 | 379.2 |
| 21:3 | 428.2 | 377.2 |
| 21:4 | 426.2 | 375.2 |
| 22:0 | 448.2 | 397.2 |
| 22:1 | 446.2 | 395.2 |
| 22:2 | 444.2 | 393.2 |
| 22:3 | 442.2 | 391.2 |
| 22:4 | 440.2 | 389.2 |
| 22:5 | 438.2 | 387.2 |
| 22:6 | 436.2 | 385.2 |

**Supplementary Table 4. Assignment of plasmenyl-PE isomeric species identified in mGluR2/mGlyR.** Assignments made from **Supplementary Fig. 6**.

| *m/z* | Lipid Identity | Isomeric Species |
|---|---|---|
| 700.53 | PE(P-34:2) | PE(P-18:1_16:1)<br>PE(P-16:0_18:2) |
| 702.54 | PE(P-34:1) | PE(P-16:0_18:1)<br>PE(P-18:1_16:0) |
| 716.52 | PE(P-35:1) | PE(P-18:0_17:1)<br>PE(P-17:1_18:0)<br>PE(P-17:0_18:1)<br>PE(P-18:1_17:0) |
| 724.53 | PE(P-36:4) | PE(P-16:0_20:4) |
| 726.54 | PE(P-36:3) | PE(P-16:0_20:3)<br>PE(P-18:2_18:1)<br>PE(P-18:1_18:2) |
| 728.56 | PE(P-36:2) | PE(P-18:1_18:1)<br>PE(P-16:0_20:2) |
| 730.57 | PE(P-36:1) | PE(P-16:0_20:1)<br>PE(P-18:0_18:1) |
| 742.54 | PE(P-37:2) | PE(P-18:1_19:1)<br>PE(P-19:1_18:1) |
| 750.54 | PE(P-38:5) | PE(P-18:1_20:4) |
| 752.56 | PE(P38:4) | PE(P-16:0_22:4)<br>PE(P-18:0_20:4) |
| 756.59 | PE(P-38:2) | PE(P-16:0_22:2)<br>PE(P-18:1_20:1)<br>PE(P-20:1_18:1)<br>PE(P-18:0_20:2) |
| 758.61 | PE(P-38:1) | PE(P-18:1_20:0)<br>PE(P-20:0_18:1)<br>PE(P-18:0_20:1)<br>PE(P-20:1_18:0) |
| 774.40 | PE(P-40:7) | PE(P-18:1_22:6) |
| 776.55 | PE(P-40:6) | PE(P-18:0_22:6)<br>PE(P-18:1_22:5) |
| 778.56 | PE(P-40:5) | PE(P-18:0_22:5)<br>PE(P-18:1_22:4) |
| 780.55 | PE(P-40:4) | PE(P-18:0_22:4) |
| 782.61 | PE(P-40:3) | PE(P-18:1_22:2)<br>PE(P-18:0_22:3) |
| 784.63 | PE(P-40:2) | PE(P-18:1_22:1)<br>PE(P-18:0_22:2) |

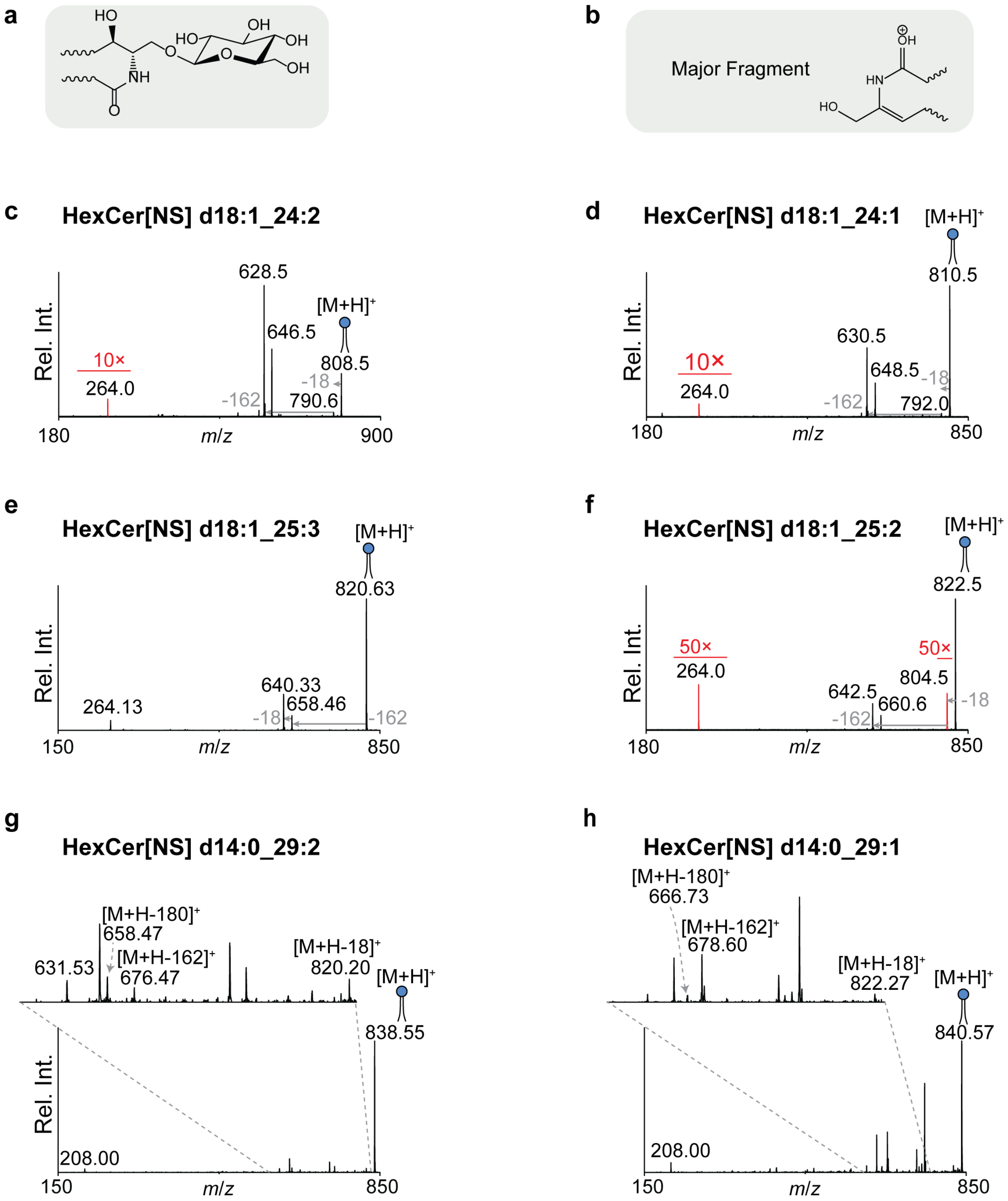

**Supplementary Figure 7. Assignment of HexCer[NS] lipids.**

**a,** Structure of HexCer[NS] headgroup. **b,** Chemical structure of common diagnostic fragment. **c-h,** Ion-trap CID fragmentation HexCer[NS] species identified in mGluR2/mGlyR. Spectra were collected as $MS^3$ from mGluR2/mGlyR incubated with brain lipids. The dominant fragments include neutral losses of water (−18 Da) and cleavage of the glycosidic bond (−162 Da). A combined loss of 180 Da is also observed. In addition, the diagnostic sphingoid base fragment at *m*/*z* 264 is observed. There are also characteristic fragments for dehydrated ceramide (*5*).

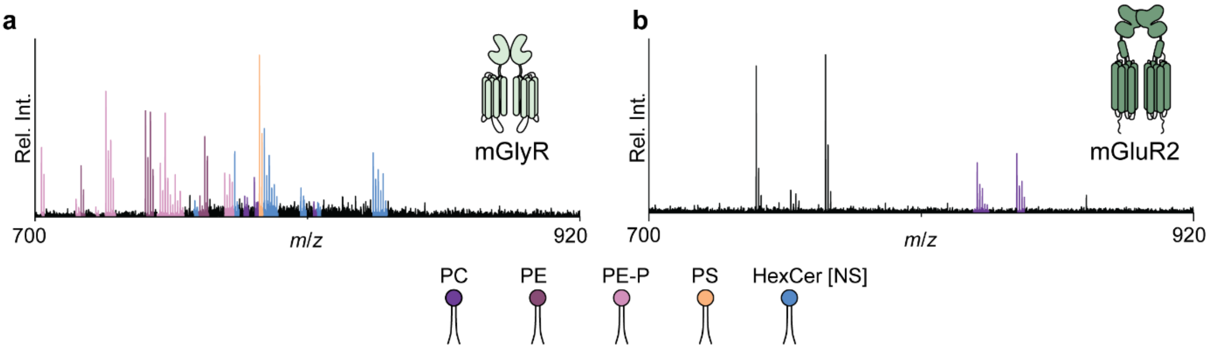

**Supplementary Figure 8. MS$^2$ of mGlyR and mGluR2 from REVEAL method.**
**a,** Representative MS$^2$ of mGlyR incubated with a ten-fold molar excess of brain lipids in the REVEAL method (quadrupole selection at *m*/*z* 6000, 10 *m*/*z* wide, 120 V HCD activation). Peaks corresponding to released lipids are seen. **b,** Representative MS$^2$ of mGluR2 incubated with a ten-fold molar excess of brain lipids in the REVEAL method (quadrupole selection at *m*/*z* 7000, 10 *m*/*z* wide, 120 V HCD activation).

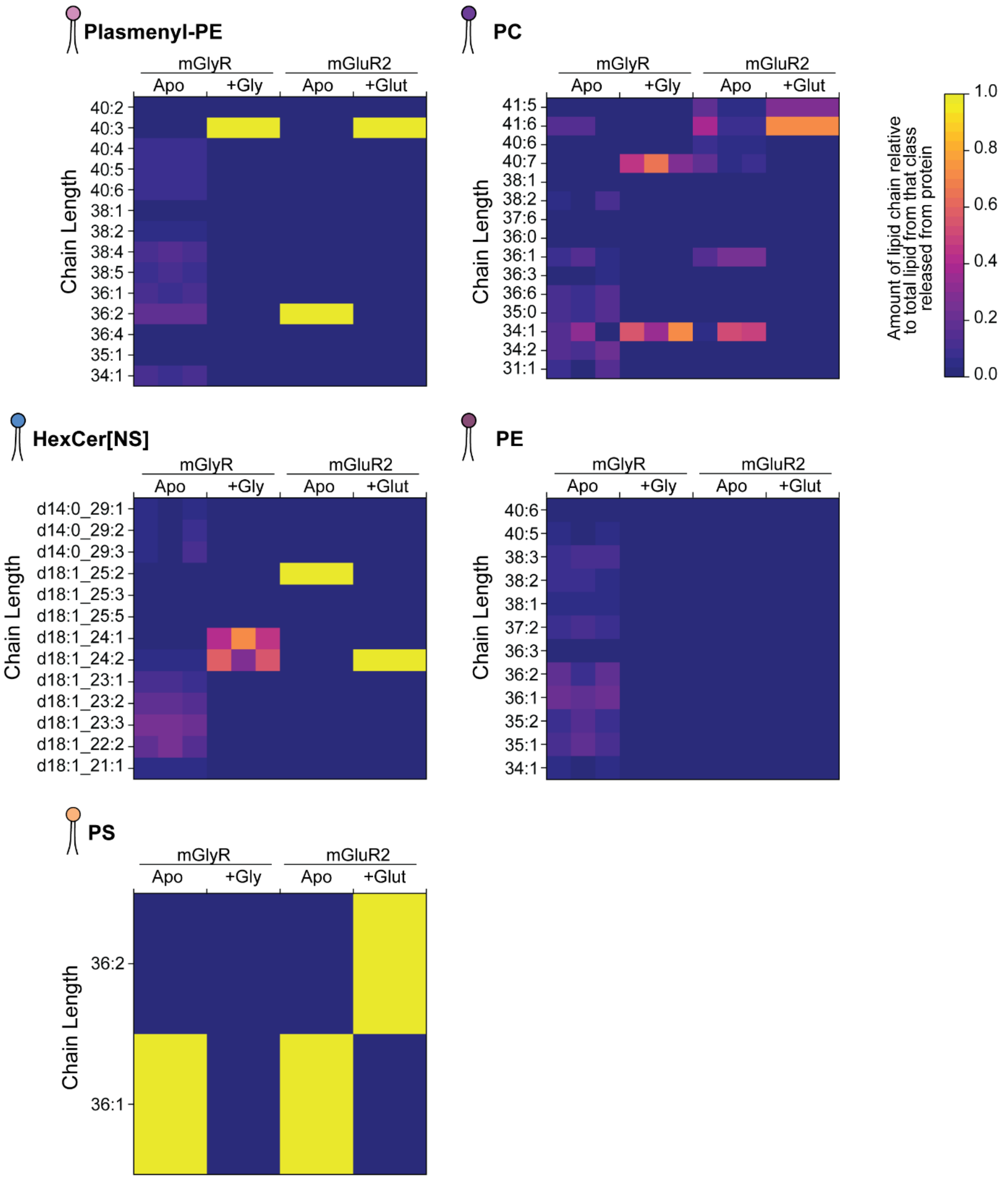

**Supplementary Figure 9. Heatmaps of lipids binding to apo and agonised mGlyR and mGluR2.** The colours represent the relative amount of each chain length as a total of that lipid class released from the protein in n=3 technical replicates. The intensities are measured from the area under the curve in the reconstructed precursor spectra from the REVEAL method.

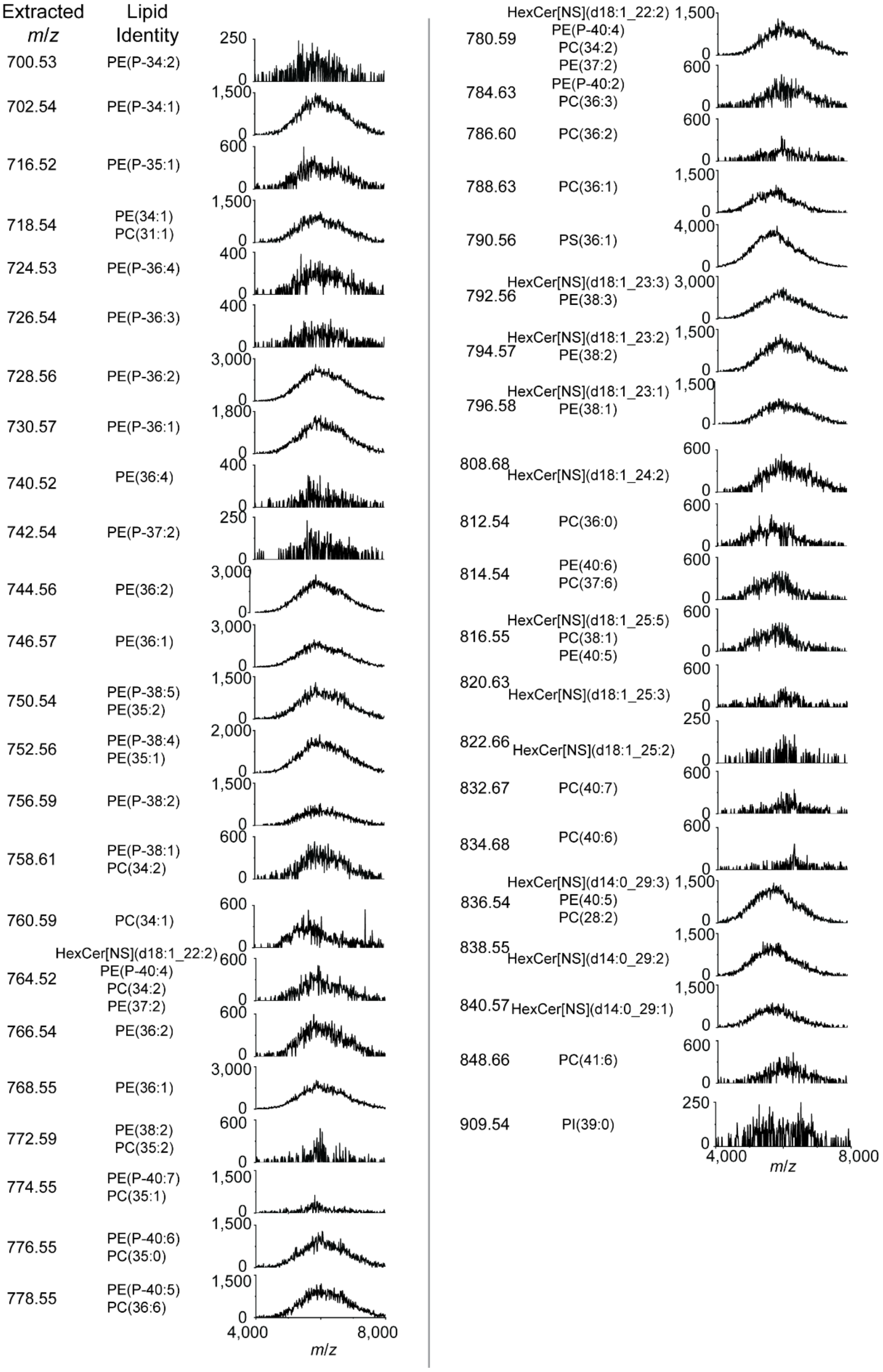

**Supplementary Figure 10. Representative reconstructed precursor spectra for all lipids in apo mGlyR.** Lipids were identified via a wide isolation nTDMS experiment, and reconstruction follows application of REVEAL. Reconstructed precursor spectra are plotted as an absolute intensity (arbitrary units).

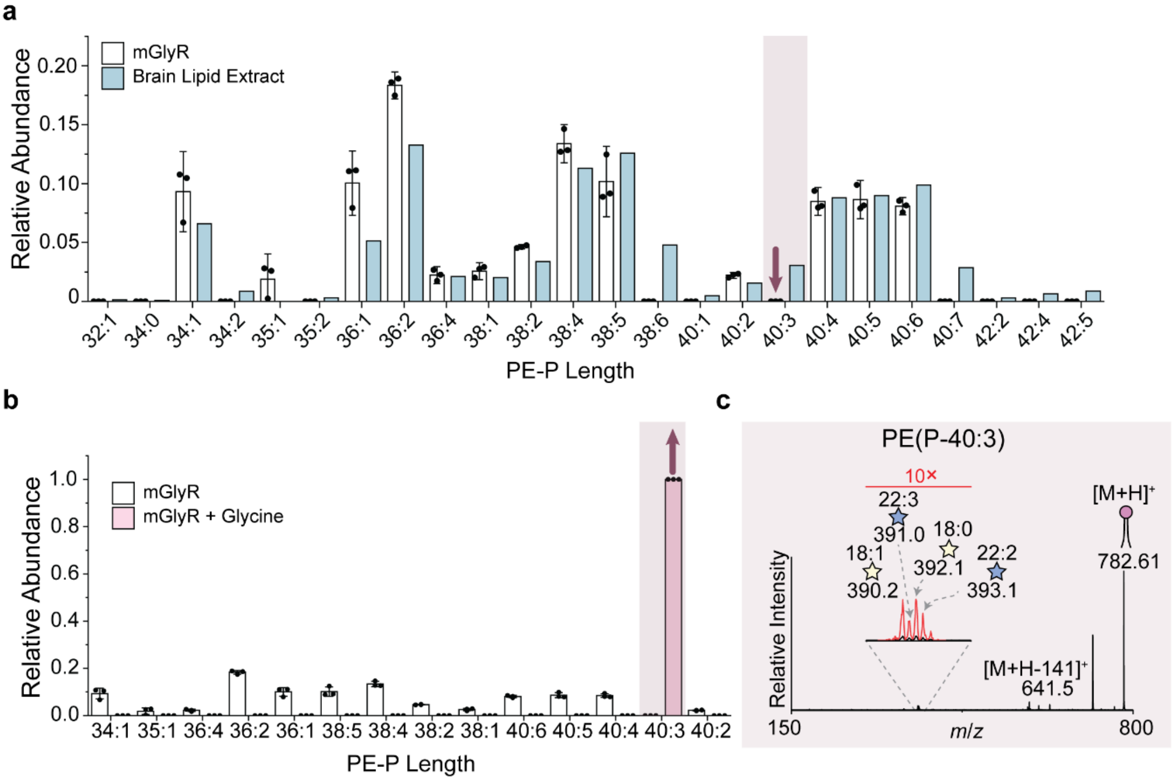

**Supplementary Figure 11. Relative abundances of PE-P lipid lengths released from mGlyR compared to the brain lipid extract.**
**a**, Bar plot showing the relative abundance of each PE-P species released from mGlyR (white), quantified by the area under the reconstructed precursor spectrum, alongside their relative abundance in brain lipid extracts measured by lipidomics (blue). Lipid abundance is expressed as a proportion of total PE-P detected. PE(P-40;3) is highlighted. **b,** Bar plot showing the relative abundance of each PE-P species released from mGlyR, with (pink) and without (white) glycine quantified by the area under the reconstructed precursor spectrum. Lipids are expressed as a proportion of total PE-P detected. PE(P-40;3) is highlighted. **c,** $MS^3$ of PE(P-40:3), as in **Supplementary Figure 6**.

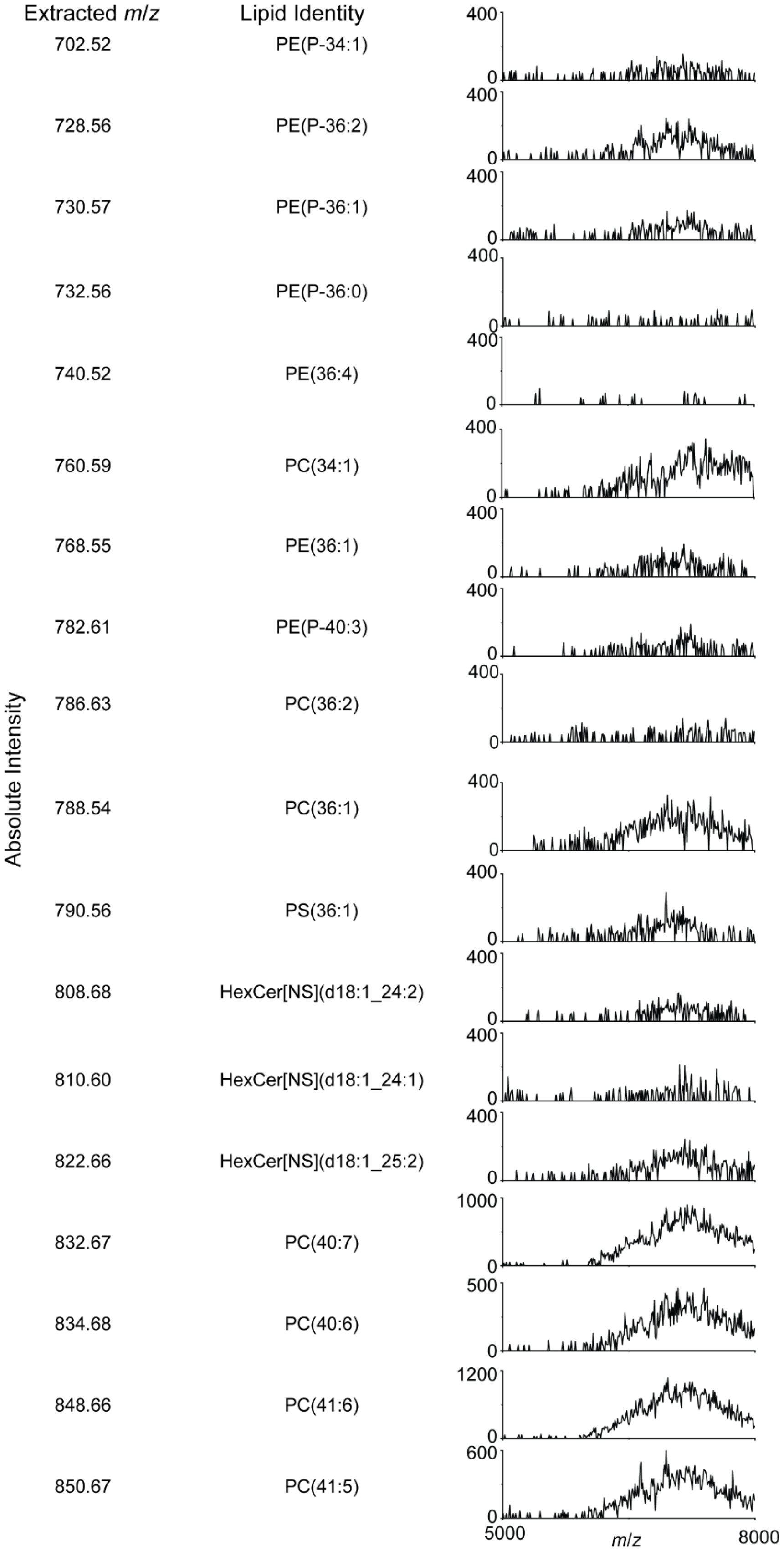

**Supplementary Figure 12. Representative reconstructed precursor spectra for all lipids in apo mGluR2.** Lipids were identified via a wide isolation nTDMS experiment, and reconstruction follows application of REVEAL. Reconstructed precursor spectra are plotted as an absolute intensity (arbitrary units).

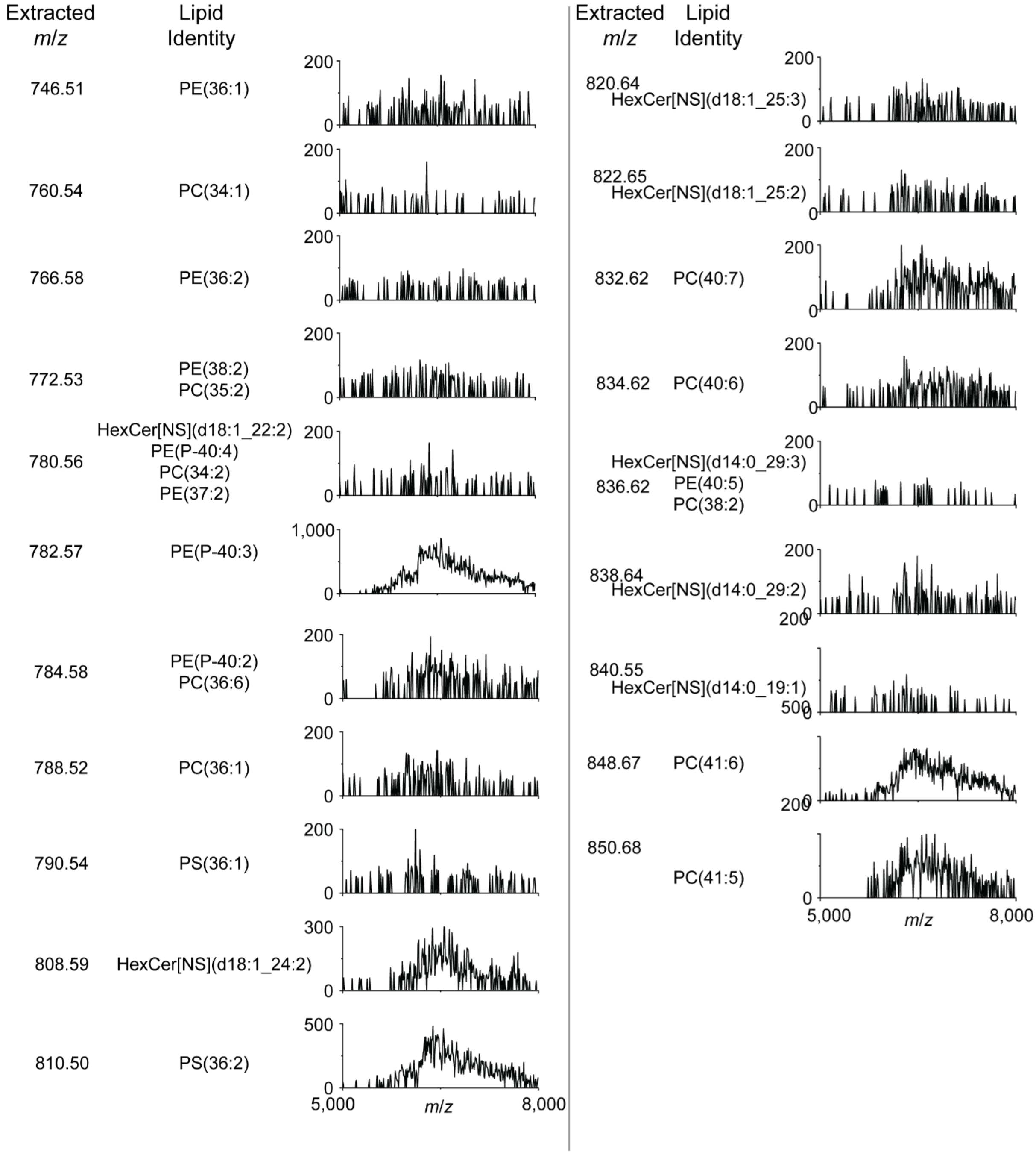

**Supplementary Figure 13. Reconstructed Precursor Spectra of glutamate-bound mGluR2.**
Lipids were identified via a wide isolation nTDMS experiment, and reconstruction following application of REVEAL. Reconstructed precursor spectra are plotted as an absolute intensity (arbitrary units).

a

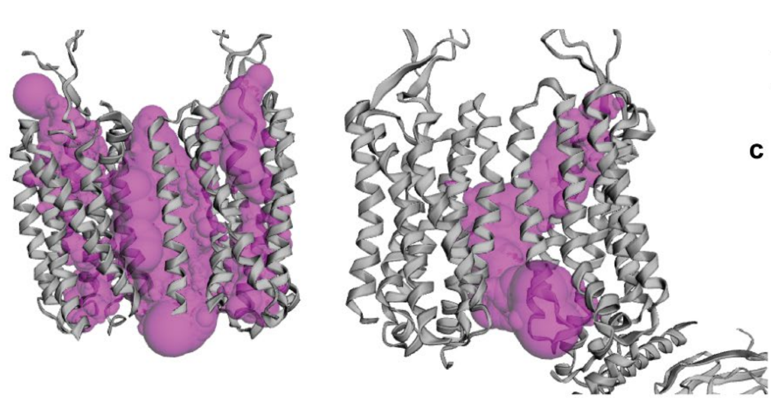

b

| | Area (Å$^2$) | Volume (Å$^3$) |
|---|---|---|
| Inactive | 6832.557 | 9150.857 |
| Active | 1953.907 | 2498.906 |

c

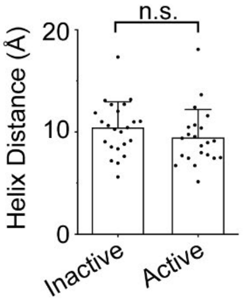

**Supplementary Figure 14. Structural Characterisation of mGluR2.**
**a**, Surface representation of active and inactive mGluR2 showing the largest predicted cavity/pocket in the TM helices identified by CASTpFold (*6*). **b,** Pocket volumes and surface areas were calculated based on the protein's folded conformation, highlighting potential ligand-binding or functional sites. **c,** Bar plot of the distance between adjacent TM alpha helices on the same protomer. Distances were measured using C$\alpha$ at the intra and extracellular end of each TM alpha helix. Distances were measured using ChimeraX. Structures used for this analysis: activated mGluR2 (PDB 7MTS), inactive mGluR2 (PDB 7MTQ).

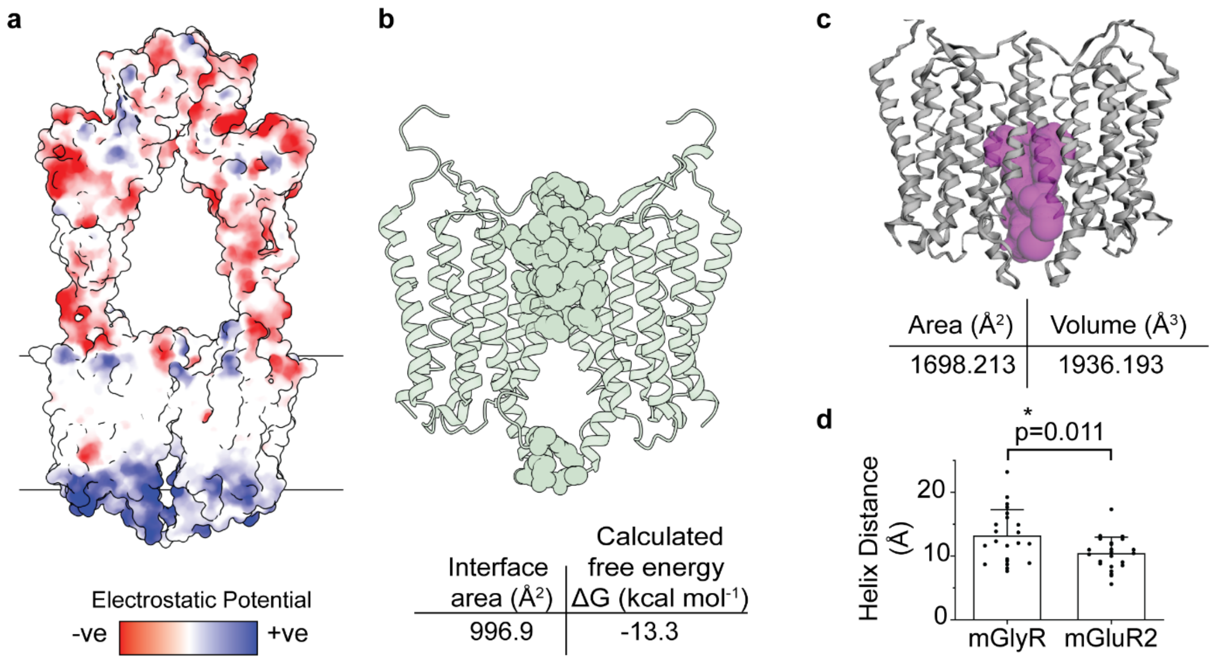

**Supplementary Figure 15. Structural Characterisation of apo mGlyR (PDB: 7SHE).**
**a,** Electrostatic surface potential of mGlyR. The red regions are more negatively charged and the blue are more positively charged. Electrostatic surfaces were modelled using ChimeraX (*7*). **b,** Interfaces between the TM helices. Residues involved in forming interactions (defined using PDBePISA) are coloured and interfacial surface areas and solvent energy gain at complex formation are listed (both calculated using PDBePISA) (***8***). **c,** Surface representation of mGlyR showing the largest predicted cavity/pocket in the TM helices identified by CASTpFold (*6*). Pocket volumes and surface areas were calculated based on the protein's folded conformation, highlighting potential ligand-binding or functional sites. **d,** Bar plot of the distance between adjacent TM alpha helices on the same protomer in mGlyR and mGluR2 (inactive, PDB 7MTQ). Distances were measured using Cα at the intra and extracellular end of each TM alpha helix. Distances were measured using ChimeraX. P value is listed.

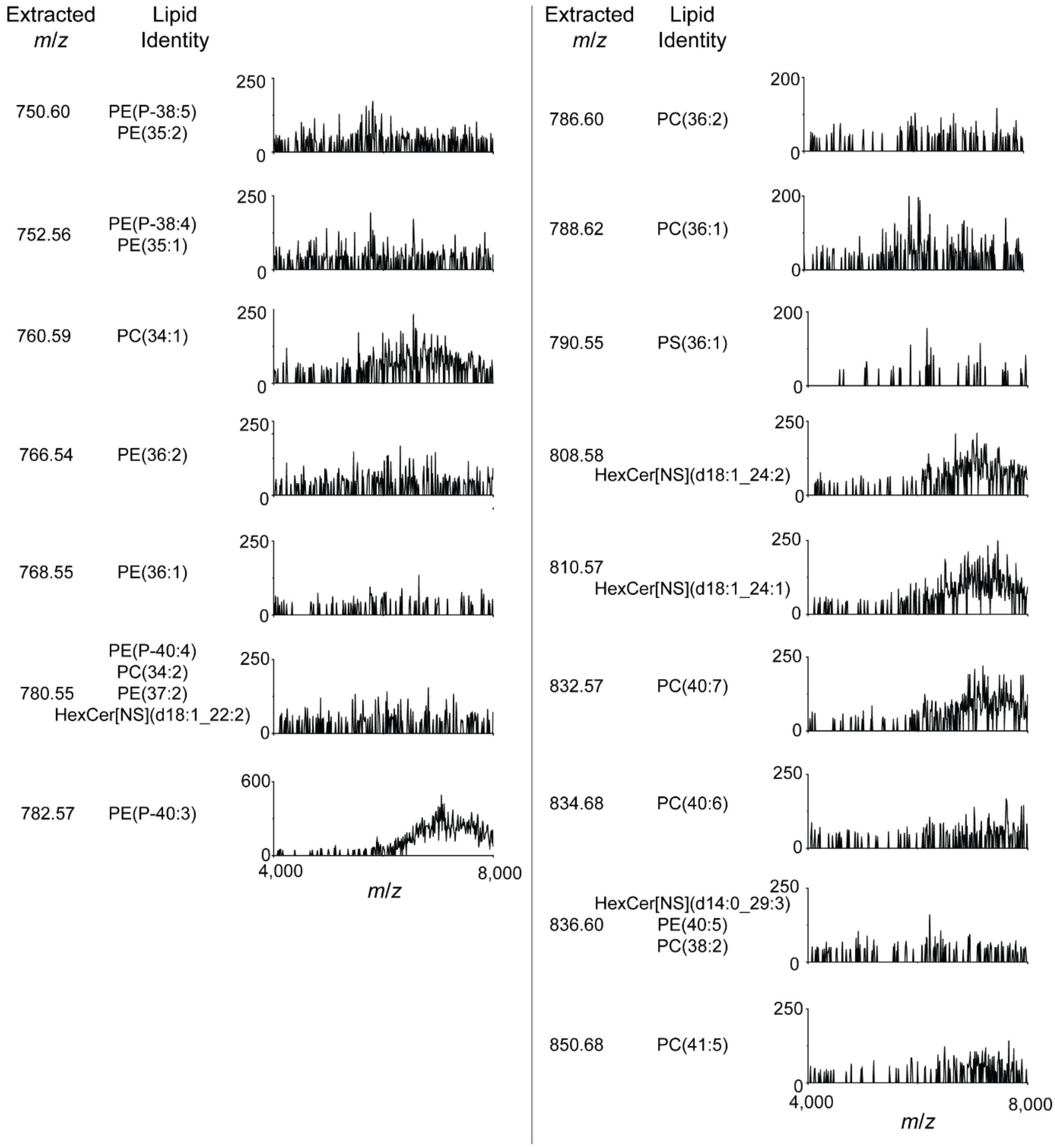

**Supplementary Figure 16. Representative reconstructed precursor spectra for lipids in activated mGlyR.** Lipids were identified via a wide isolation targeted nTDMS experiment, and reconstruction following application of REVEAL. Reconstructed precursor spectra are plotted as an absolute intensity (arbitrary units).

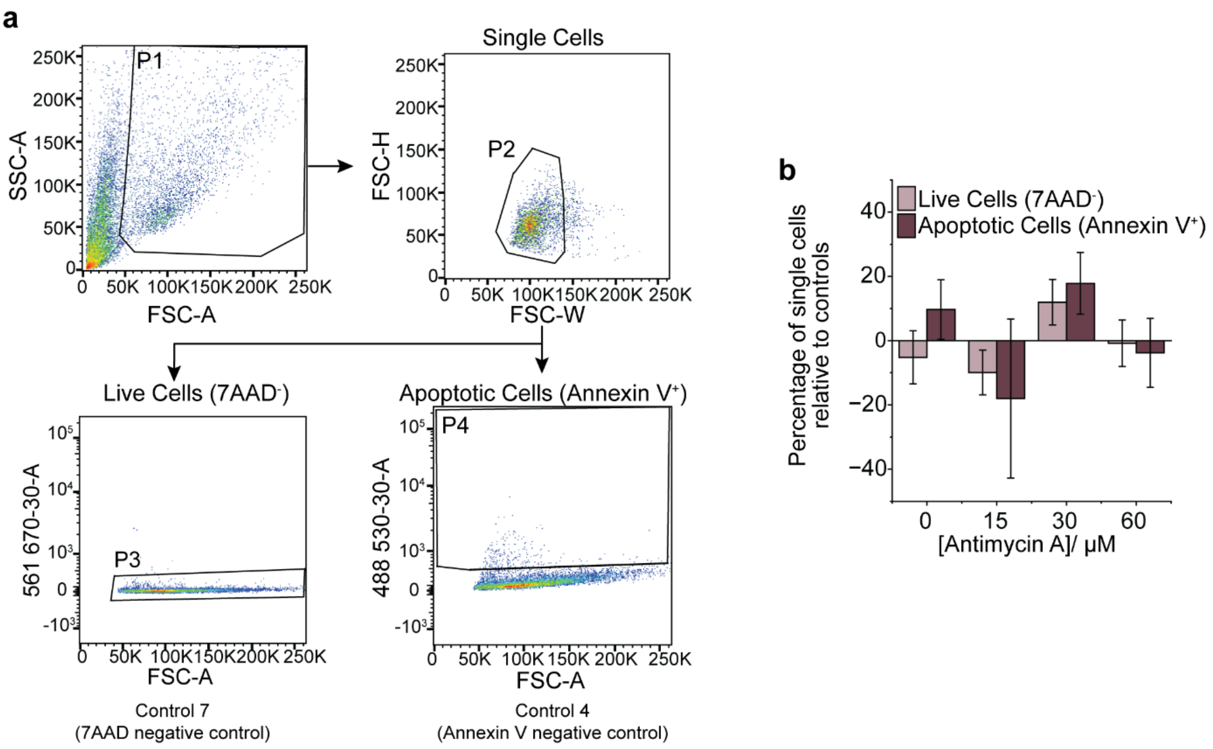

**Supplementary Figure 17. Flow cytometric analysis of cell viability and apoptosis in HEK293T cells following Antimycin A treatment.**

**a,** Gating strategy for HEK293T cells stained with Annexin V and 7-AAD. Single cells were first identified and subsequently gated to distinguish viable (Annexin $V^{-}$/7-$AAD^{-}$), early apoptotic (Annexin $V^{+}$/7-$AAD^{-}$), and late apoptotic or dead (7-$AAD^{+}$) populations. **b,** Quantification of the percentage of single cells that are 7-$AAD^{-}$ and Annexin $V^{+}$ at increasing drug concentrations, normalised to untreated control cells.

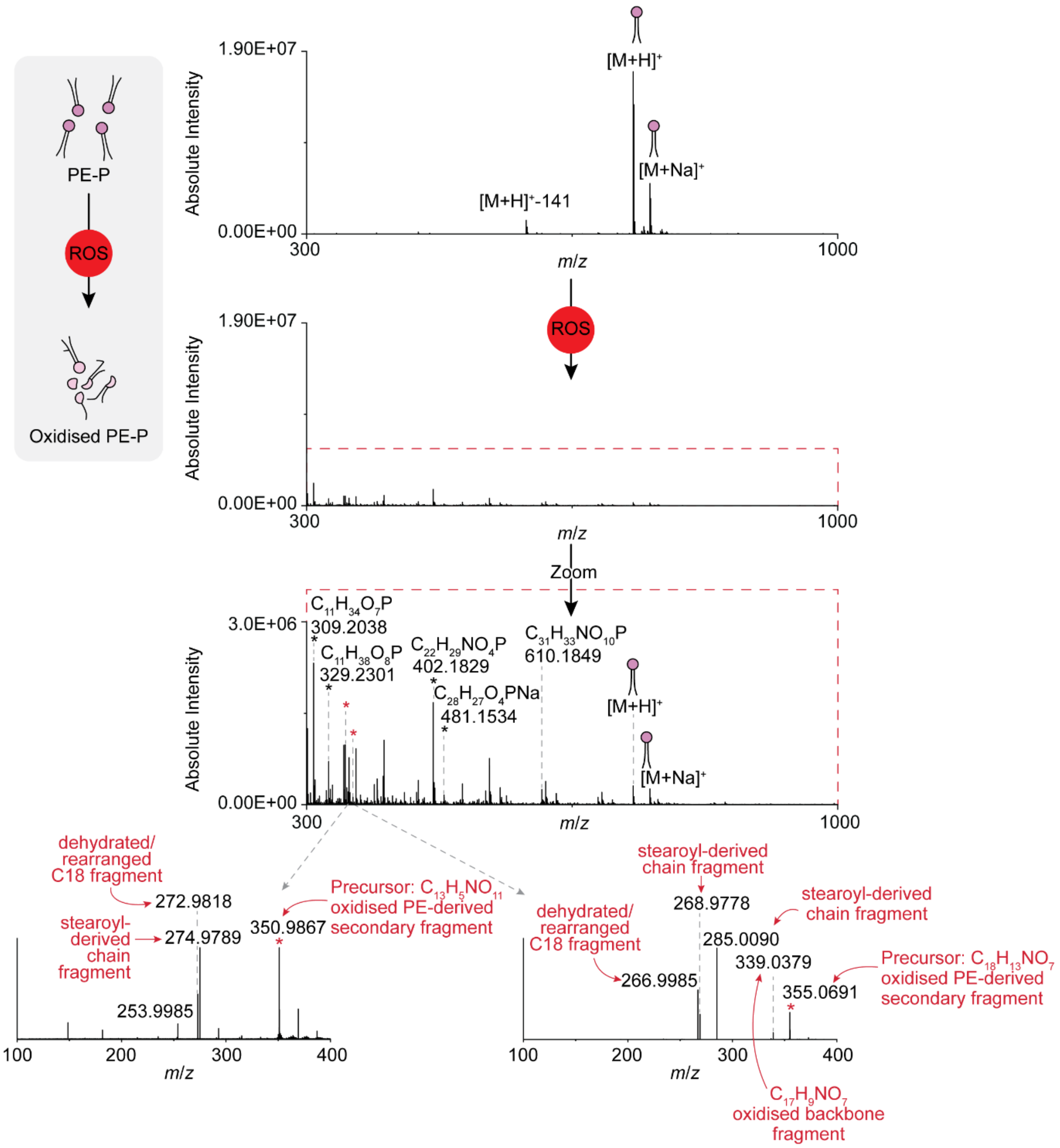

**Supplementary Figure 18. Mass spectrometric analysis of PE(P-18:0/18:1) following radical oxidation.** Positive-ion mass spectra of a PE(P-18:0/18:1) standard acquired before and after induction of radical oxidation using $H_2O_2$ and $Fe^{2+}$ (Fenton conditions). The untreated sample shows protonated ($[M+H]^+$) and sodiated ($[M+Na]^+$) ions corresponding to the intact plasmalogen, along with a characteristic headgroup loss ($[M+H]^+$ − 141 Da). Following oxidation, new product ions are observed, consistent with radical-mediated modification and fragmentation of the plasmalogen backbone. A zoomed region highlights representative oxidation-derived ions, including truncated and oxidised phospholipid species. Selected ions were subjected to $MS^2$ analysis to support chemical assignment. Fragmentation spectra reveal characteristic C18 chain-derived ions and oxidised backbone fragments, consistent with cleavage at the glycerophospholipid backbone. Peaks were assigned chemical formula based on accurate mass (≤3 ppm error) and fragmentation behaviour.

**Supplementary Table 5. Adjusted P values accompanying Fig. 4i.**

| Group 1 | Group 2 | Mean Difference | Adjusted P | Lower CI | Upper CI | Reject Null | Significance Level |
|---|---|---|---|---|---|---|---|
| CONTROL (1) | CONTROL +AA (2) | -0.4852 | 0.0023 | -0.7886 | -0.1818 | TRUE | ** |
| CONTROL (1) | mGlyR (3) | -0.285 | 0.068 | -0.5884 | 0.0184 | FALSE | ns |
| CONTROL (1) | mGlyR +AA (4) | -0.8113 | 0 | -1.1147 | -0.5079 | TRUE | *** |
| CONTROL +AA (2) | mGlyR (3) | 0.2002 | 0.2561 | -0.1033 | 0.5036 | FALSE | ns |
| CONTROL +AA (2) | mGlyR +AA (4) | -0.3261 | 0.034 | -0.6295 | -0.0227 | TRUE | * |
| mGlyR (3) | mGlyR +AA (4) | -0.5263 | 0.0012 | -0.8297 | -0.2229 | TRUE | ** |

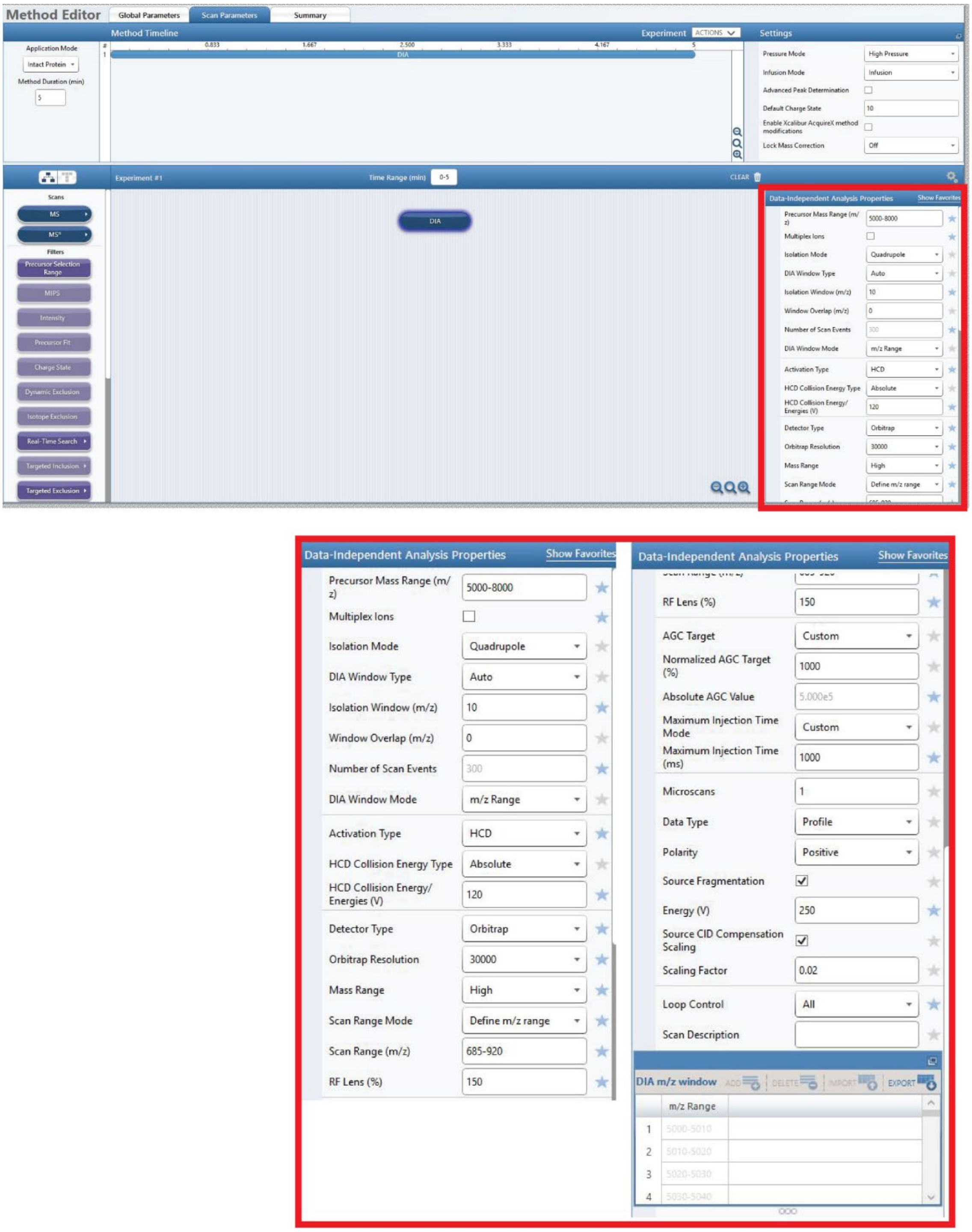

**Supplementary Figure 19: Example REVEAL method used on a Thermo Fisher Orbitrap Ascend Tribrid mass spectrometer.**
Method was written in XCalibur (version 4.6.67.17).
